# Tudor staphylococcal nuclease acts as a docking platform for stress granule components in *Arabidopsis thaliana*

**DOI:** 10.1101/2020.02.20.955922

**Authors:** Emilio Gutierrez-Beltran, Pernilla H. Elander, Kerstin Dalman, Jose Luis Crespo, Panagiotis N. Moschou, Vladimir N. Uversky, Peter V. Bozhkov

**Affiliations:** Instituto de Bioquímica Vegetal y Fotosíntesis, Universidad de Sevilla and Consejo Superior de Investigaciones Científicas, Sevilla, Spain; Department of Molecular Sciences, Uppsala BioCenter, Swedish University of Agricultural Sciences and Linnean Center for Plant Biology, PO Box 7015, SE-75007 Uppsala, Sweden; Institute of Molecular Biology and Biotechnology, Foundation for Research and Technology - Hellas, Heraklion, Greece; Department of Plant Biology, Uppsala BioCenter, Swedish University of Agricultural Sciences and Linnean Center for Plant Biology, PO Box 7080, SE-75007 Uppsala, Sweden; Department of Biology, University of Crete, Heraklion, Greece; Department of Molecular Medicine, Morsani College of Medicine and USF Health Byrd Alzheimer’s Research Institute, University of South Florida, Tampa, FL, USA; Institute for Biological Instrumentation of the Russian Academy of Sciences, Federal Research Center “Pushchino Scientific Center for Biological Research of the Russian Academy of Sciences”, Pushchino, Moscow region, 142290, Russia

**Keywords:** intrinsically disordered region (IDR), liquid-liquid phase separation (LLPS), messenger ribonucleoprotein (mRNP) complex, RNA-binding protein (RBP), SG proteome, stress granule (SG), SNF1-related protein kinase 1 (SnRK1), Tudor Staphylococcal Nuclease (TSN).

## Abstract

Adaptation to stress depends on the modulation of gene expression. Regulation of mRNA stability and degradation in stress granules (SGs), - cytoplasmic membraneless organelles composed of messenger ribonucleoprotein (mRNP) complexes, - plays an important role in fine-tuning of gene expression. In addition, SG formation can modulate stress signaling pathways by protein sequestration. Molecular composition, structure, and function of SGs in plants remain obscure. Recently, we established Tudor Staphylococcal Nuclease (TSN or Tudor-SN; also known as SND1) as integral component of SGs in *Arabidopsis thaliana*. Here, we combined purification of TSN interactome with cell biology, reverse genetics and bioinformatics to study composition and function of SGs in plants. We found that under both normal (in the absence of stress) and stress conditions TSN interactome is enriched in the homologues of known mammalian and yeast SG proteins, in addition to novel or plant-specific SG components. We estimate that upon stress perception, approximately half of TSN interactors are recruited to SGs *de novo*, in a stress-dependent manner, while another half represent a dense protein-protein interaction network pre-formed before onset of stress. Almost all TSN-interacting proteins are moderately or highly disordered and approximately 20% of them are predisposed for liquid-liquid phase separation (LLPS). This suggests that plant SGs, similarly to mammalian and yeast counterparts, are multicomponent viscous liquid droplets. Finally, we have discovered that evolutionary conserved SNF1-related protein kinase 1 (SnRK1) interacts with TSN in heat-induced SGs and that SnRK1 activation critically depends on the presence of TSN and formation of SGs. Altogether, our results establish TSN as a docking platform for SG-associated proteins and important stress signal mediator in plants.

## INTRODUCTION

Upon stress perception, eukaryotic cells compartmentalize specific mRNA molecules stalled in translation initiation in two types of evolutionarily conserved membraneless organelles (MLOs) called stress granules (SGs) and processing bodies (PBs) (Thomas et al., 2011; Protter and Parker, 2016). In these organelles, mRNA molecules are stored, degraded or kept silent to prevent energy expenditure on producing useless, surplus or even harmful proteins under stress conditions.

Recent research in budding yeast *Saccharomyces cerevisiae* and animal models established the molecular composition of SGs and PBs. SGs typically contain poly(A)+ mRNA, 40S ribosomal subunits, various translation initiation factors (eIF), poly(A)-binding protein (PABP) and a variety of RNA-binding proteins (RBPs) and non-RNA-binding proteins (Buchan and Parker, 2009). PBs contain proteins belonging to the mRNA decay machinery, including subunits of decapping and exosome complexes (DCP and XRN proteins, respectively), deadenylases and many RBPs (Franks and Lykke-Andersen, 2008). Although there is a dynamic flux of proteins and mRNA molecules between SGs and PBs, these MLOs have different functions. SGs play a major role in translational repression by sequestering, stabilizing and storing mRNA molecules, as well as by indirectly modulating signaling pathways, whereas PBs are to a large extent involved in mRNA decay (Protter and Parker, 2016; Mahboubi and Stochaj, 2017).

Apart from components of SGs and PBs, proteomic and genetic screens in yeast and animal models have also identified proteins modulating their assembly, which is a highly coordinated process driven by numerous proteins (Ohn et al., 2008; Buchan et al., 2011; Martinez et al., 2013; Jain et al., 2016). A recent model for the formation of mammalian and yeast SGs suggest that they assemble in a two-step process, first involving the formation of a dense stable SG core followed by accumulation of proteins containing intrinsically disordered regions (IDRs) into a peripheral shell, the process involving liquid-liquid phase separation (LLPS) (Jain et al., 2016; Markmiller et al., 2018).

In plants, molecular composition and function of SGs, as well as pathways of their assembly and cross-talk with other signaling pathways remain largely unknown. Previous studies in Arabidopsis thaliana (*Arabidopsis*) revealed formation of SGs under heat, hypoxia and salt stress (Sorenson and Bailey-Serres, 2014; Yan et al., 2014; Gutierrez-Beltran et al., 2015). To date, few conserved components of plants SGs have been identified, most of which are homologues of yeast and/or mammalian SG-associated proteins. These include T-cell-restricted intracellular antigen-1 (TIA-1) homologues Rbp47 and oligouridylate-binding protein 1 (Ubp1), endoribonuclease G3BP homologue nuclear transport factor 2 (NTF2-RPM) and tandem CCCH zinc finger (TZF) family of RBPs (Bogamuwa and Jang, 2013; Sorenson and Bailey-Serres, 2014; Gutierrez-Beltran et al., 2015; Krapp et al., 2017; Kosmacz et al., 2018).

Interestingly, Tudor Staphylococcal Nuclease (TSN) has been established as a novel component of SGs in such distant lineages as protozoa, animals and plants (Zhu et al., 2013; Yan et al., 2014; Gao et al., 2015; Gutierrez-Beltran et al., 2015; Cazares-Apatiga et al., 2017). The domain composition of TSN includes tandem repeat of four Staphylococcal Nuclease (SN) domains at its N-terminus followed by a Tudor and C-terminal, partial SN domain (Abe et al., 2003; Gutierrez-Beltran et al., 2016). To date, TSN is known to be critically involved in the regulation of virtually all pathways of gene expression, ranging from transcription to RNA silencing (Gutierrez-Beltran et al., 2016). The interaction of TSN with proteins forming the core of SGs, such as Pabp1, eIF4E, eIF5A, TIAR (TIA1-related protein) and TIA1 (T-cell-restricted intracellular antigen) in different organisms indicate that TSN is an evolutionary conserved component of SGs (Weissbach and Scadden, 2012; Zhu et al., 2013; Gao et al., 2014).

In yeast and mammals, the universal molecular components of SGs co-exist together with other cell type- and stress stimulus-specific components, suggesting that SGs might play additional, yet unexplored, roles during stress. Thus, SG assembly in both yeast and human cells affects target of rapamycin (TOR) signaling by sequestering both complex 1 TOR and downstream kinases to alter signaling during stress (Takahara and Maeda, 2012; Wippich et al., 2013). On the other hand, sequestration of the pleiotropic adaptor protein Receptor For Activated C Kinase 1 (RACK1) in SGs inhibits the stress-induced activation of the c-Jun N-terminal kinases (JNK) cascade that triggers apoptotic death (Arimoto et al., 2008). In another scenario, sequestration of the coiled-coil containing protein kinase 1 (ROCK1) into SGs promotes cell survival by abolishing JNK-mediated cell death (Tsai and Wei, 2010). In summary, SG formation can control signaling pathways by protein sequestration during stress conditions, but whether such mode of regulation exists in plants remains elusive.

In the present study, we isolated TSN-interacting proteins from *Arabidopsis* plants subjected to different types of stresses, and further combined microscopy, reverse genetics and bioinformatics to advance our understanding of the regulation and molecular function of SGs in plants. We show that SG proteins form a dense interaction network already under normal (no stress) conditions that is poised to enable rapid SG assembly in response to stress. We found that TSN functions as a platform for docking homologues of key components of yeast and mammalian SGs, as well as novel or plant-specific components. Ubiquitous occurrence of intrinsically-disordered proteins (IDPs) among TSN interactors supports a notion of LLPS being a key process underlying SG assembly in plants. Finally, we have discovered that TSN and formation of SGs confer heat-induced activation of the major energy sensor SNF1-related protein kinase 1 (SnRK1).

## RESULTS

### Generation and characterization of Arabidopsis TAPa-expressing lines

As a first step to investigate the role of TSN in SG formation, we have used TSN2 *Arabidopsis* isoform as a bait for alternative tandem affinity purification (TAPa) (Rubio et al., 2005). Thus, TSN2 and green fluorescent protein (GFP; negative control) were tagged at their C-termini with TAPa epitope containing two copies of the immunoglobulin-binding domain of protein A from *Staphylococcus aureus*), a human rhinovirus 3 protease cleavage site, a 6-histidine repeat and 9-myc epitopes (Figure 1A). Resulting TSN2-TAPa and GFP-TAPa vectors were introduced into *Arabidopsis* Columbia (Col) background. Two lines per construct showing good expression levels were selected for further studies (Supplemental Figure S1A).

**Figure 1.**
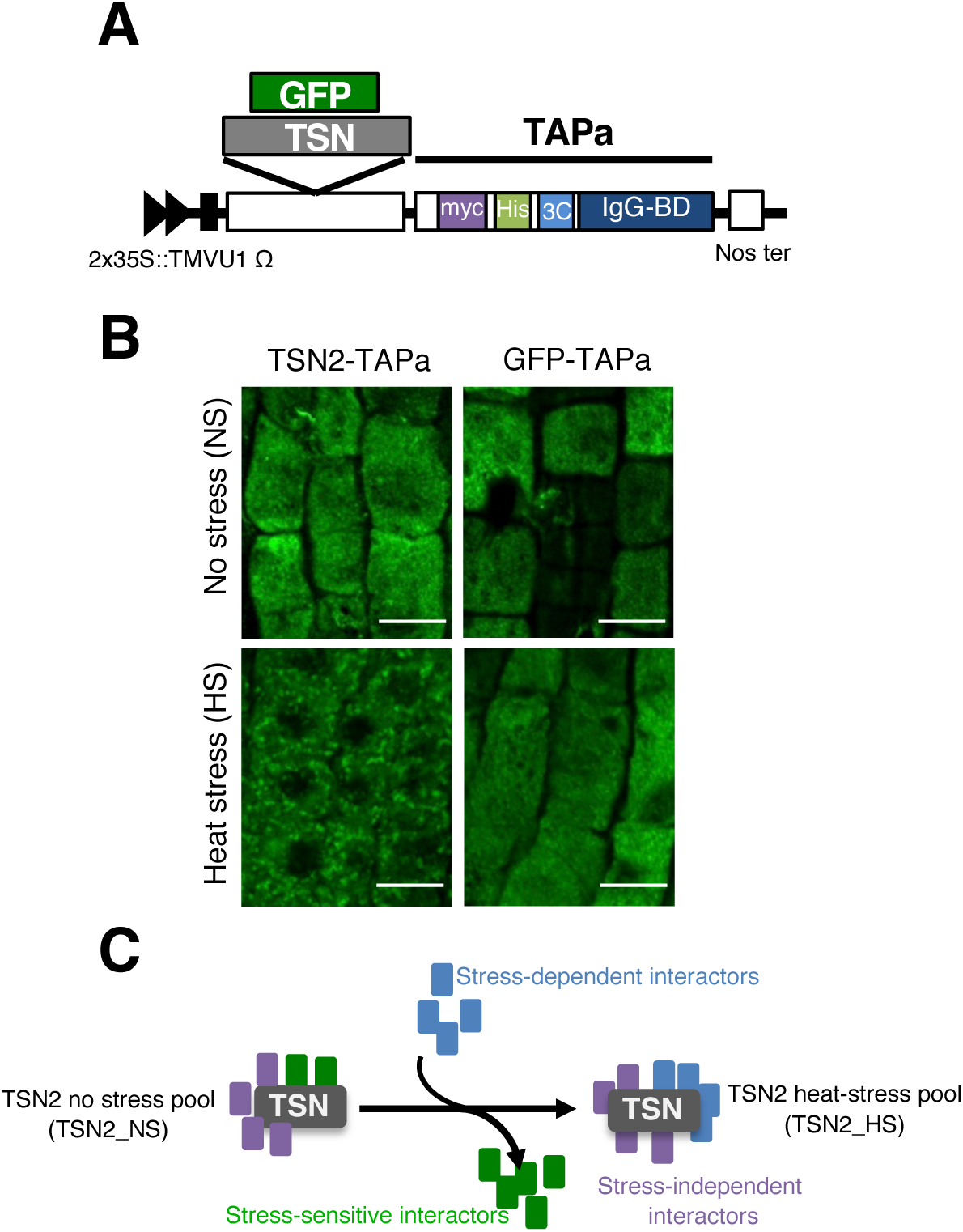
Identification of *Arabidopsis* TSN-interacting proteins by alternative tandem affinity purification (TAPa). ***A***, Schematic illustration of the expression cassette in TAPa vector. The vector allows translational fusion of TSN and GFP at their C-termini to the TAPa tag. The expression is driven by two copies of the cauliflower mosaic virus 35S promoter (2×35S) and a tobacco mosaic virus (TMV) U1 X translational enhancer. The TAPa tag consists of two copies of the protein A IgG-binding domain (IgG-BD), an eight amino acid sequence corresponding to the 3C protease cleavage site (3C), a 6-histidine stretch (His), and nine repeats of the Myc epitope (myc). A Nos terminator (Nos ter) sequence is located downstream of each expression cassette. ***B***, Immunolocalization of TSN2-TAPa and GFP-TAPa fusion proteins in root cells of 5-day-old seedling. The seedlings were grown under no stress conditions (23°C) or incubated for 40 min at 39°C (heat stress) and then immunostained with α-Myc. Scale bars = 10 µm. ***C***, Schematic representation of three classes of proteins found upon comparative analysis of TSN interactomes isolated under no stress and heat stress conditions.

To verify whether the presence of TAPa epitope could affect intracellular localization of TSN protein, we performed immunostaining of control and heat-stressed root cells using α-Myc. The analysis revealed that similar to native TSN (Yan et al., 2014; Gutierrez-Beltran et al., 2015), TSN2-TAPa had diffused cytoplasmic localization under no stress (NS) conditions but redistributed to punctate foci following heat stress (HS) (Figure 1B). In contrast, GFP-TAPa remained cytoplasmic regardless of conditions (Figure 1B). These data suggested that C-terminally TAPa-tagged TSN seemed to preserve key functional features of its native counterpart when expressed in *Arabidopsis*, thus representing physiologically-relevant bait for the isolation of TSN-interacting proteins.

### Identification of TSN2-interacting proteins

To examine the feasibility and efficiency of purifying TSN2-TAPa and GFP-TAPa proteins from the corresponding transgenic plants, we performed a small-scale TAPa purification by following the TAPa purification procedure (Figure S2). As shown in Figure S3A, the immunoblot analysis with α-Myc confirmed that both TAPa-tagged proteins could be properly purified. Since TSN2 was originally identified as a robust marker of SGs induced by HS in *Arabidopsis* plants (Gutierrez-Beltran et al., 2015), we anticipated to initially compare the TSN2 interactomes under HS and NS conditions. This comparison enabled classification of various TSN interactors into one of three classes (Figure 1C): (i) stress-independent interactors, which associate with TSN independently of HS; (ii) stress-dependent interactors, which associate with TSN only under HS; and (iii) stress-sensitive interactors, whose association with TSN is lost during HS.

The mass spectrometry analysis yielded 1,091 and 4,490 hits under NS conditions and 1,493 and 1,573 hits under HS conditions in GFP and TSN2 isolations, respectively (Figure S4). In order to identify specific interactors of TSN2, we filtered the results using a two-step procedure. First, we removed GFP-interacting proteins as well as proteins that are not present in at least two biological replicates (non-specific interactions in Figure S4). The only exception was made for some of the well-known plant SG components, such as PAB4, Rbp47 or RHM, which were kept in the list regardless of their appearance in the GFP sample. Thereafter, proteins were filtered based on subcellular localization according to The *Arabidopsis* Subcellular Database (Tanz et al., 2013), excluding proteins found in the chloroplasts or mitochondria (subcellular localization in Figure S4). As a result, we obtained the lists of 315 and 176 proteins representing physiologically relevant interactomes of TSN2 under NS and HS settings, respectively (Figure S4 and Supplemental Table 1)

A comparative analysis of KEGG orthologs [KO; (Nakaya et al., 2013)] revealed that ∼ 28% proteins from both TSN2_NS and TSN2_HS pools are known components of human or yeast SGs (Figure 2A and Supplemental Figure S5A) (Jain et al., 2016). Similarly, subsets of 20 or 21 proteins from either pool of TSN2 interactors were shared with the recently reported *Arabidopsis* Rbpb47b interactome (Figure S5B) (Kosmacz et al., 2019). An *in silico* analysis found a significant degree of similarity among TSN2_NS pool, TSN_HS pool and both yeast and human SG proteomes in regards to functional distribution of composite proteins. Thus, both TSN2_NS and TSN2_HS pools are enriched in RNA-binding proteins (RBPs), proteins with predicted prion-like domains (PrLDs) and proteins with ATPase activity (Figure 2B, Supplemental Table 2). We could not find known PB components such as DCP or XRN family proteins among TSN2 interactors, indicating that TSN specifically binds SG components (Maldonado-Bonilla, 2014; Youn et al., 2018). Collectively these data demonstrate the robustness of our approach, in which TSN2 was chosen as bait for identification of plant SG-associated components.

**Figure 2.**
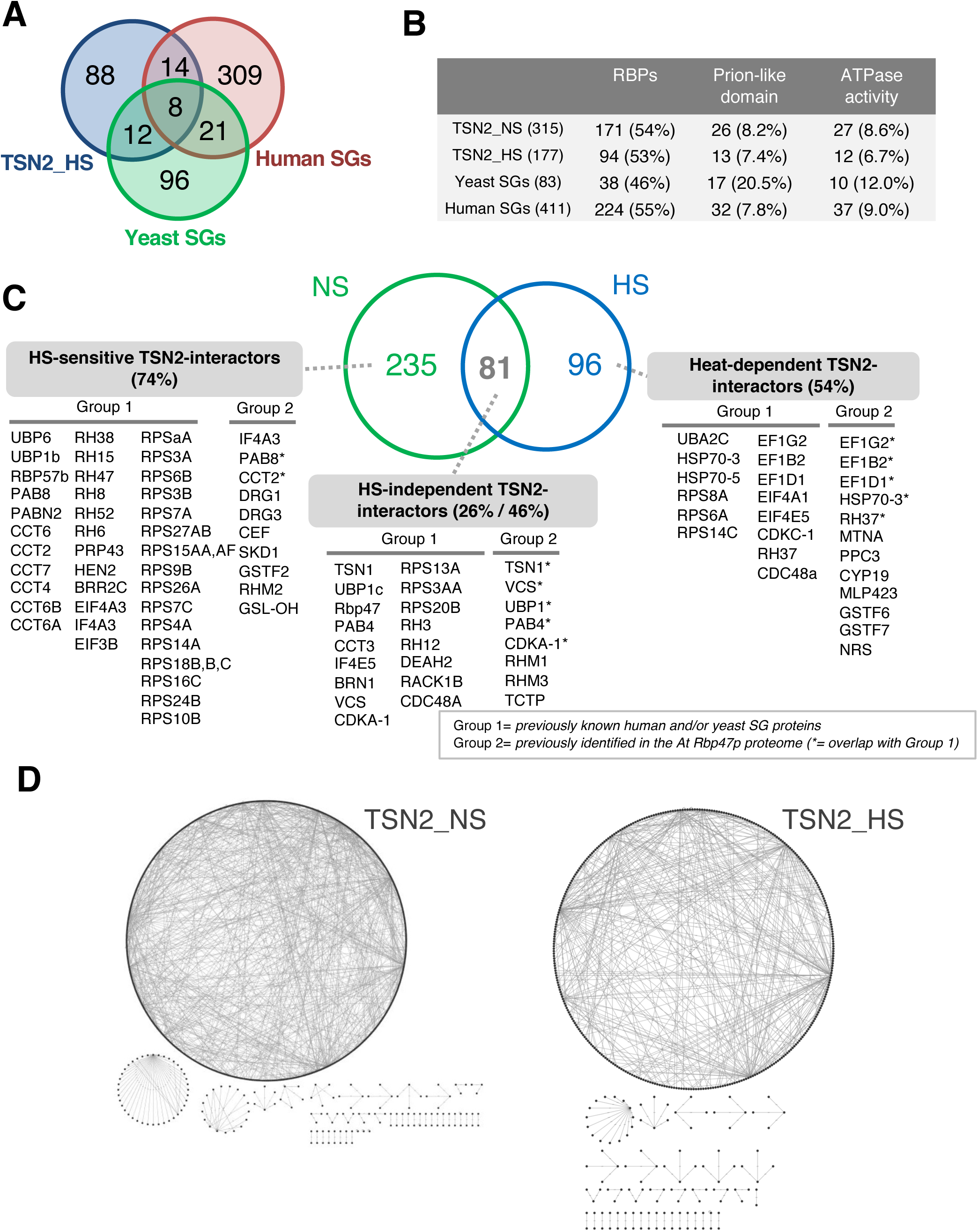
Proteomic analysis of the *Arabidopsis* TSN2 interactomes. ***A***, Venn diagram showing comparison of TSN2_HS interactome with human and yeast SG proteomes. ***B***, Frequency of RBPs and proteins with prion-like domain or ATPase activity found among TSN2_NS and TSN2_HS protein pools in comparison with yeast and human SG proteomes. ***C***, Venn diagram showing comparison between TSN2_NS and TSN2_HS protein pools. Classification of TSN2-interacting proteins in three classes: HS-sensitive, HS-independent and HS-dependent. Within each class the proteins are further classified into two groups, based on their known localization to SGs in non-plant (Group 1) or plant (Group 2) organisms. ***D***, Network maps showing physical interactions among proteins in the TSN2_NS and TSN2_HS pools.

### TSN interactome reveals a pre-existing network of SG protein-protein interactions

Although SGs are only microscopically visible under stress conditions (Jain et al., 2016), analysis of the TSN2_NS and the TSN_HS protein pools revealed that both pools are significantly enriched in known mammalian and/or yeast SG components (Group 1, Figure 2C), and also in proteins found in the previously characterized *Arabidopsis* Rbp47b proteome (Kosmacz et al., 2019), such as PAB8, SKD1, IF4A3 or RHM2 (Group 2, Figure 2C). Furthermore, 74% (235/315) protein hits of the TSN2_NS pool were absent from the TSN2_HS pool, thus representing HS-sensitive part of the TSN2 interactome. Accordingly, the remaining, smaller part of the TSN2_NS pool (26%, 81/315) was shared with the TSN2_HS pool, corresponding to stress-independent TSN2 interactors (Figure 2C). The latter class of proteins included TSN1, UBP1c, Rbp47, PAB4, VCS or TCTP, among others. Lastly, 54% (96/177) protein hits from the TSN2_HS pool, including HSP70 and individual subunits of both EIF4 translation initiation factors and EF1elongation factors were absent from the TSN2_NS pool, and therefore represented HS-dependent TSN2 interactors (Figure 2C).

To expand on these findings, we also retrieved publicly available direct protein-protein interaction (PPI) data for all proteins found in our proteomic studies. Both TSN2_NS and TSN2_HS pools formed a dense network of protein-protein interactions (Figure 2D), containing 315 and 176 nodes and 885 and 469 edges respectively. In this context, the average number of interactions per protein for these two pools was 5.4 (p = 5.63 × 10^-11^) and 4.1 (p = 2.17 × 10^-10^), respectively. Together with our previous observations that the core SG proteins such as Rbp47, Ubp1 or PAB4 interact with TSN in *Arabidopsis* cells in the absence of stress, these results point to a pre-existing steady-state network of protein interactions as a primordial mechanism during SG formation, where TSN could act as an assembly platform.

### TSN interactome is enriched in IDPs

Studies in mammalian and yeast cells have suggested that SGs are multicomponent viscous liquid droplets formed in the cytoplasm by LLPS (Kroschwald et al., 2015; Protter and Parker, 2016). Although the molecular details underlying LLPS in cells are largely obscure, recent evidence suggests that IDRs mediate this process (Molliex et al., 2015; Wheeler et al., 2016). In this context, we first evaluated the occurrence of proteins with IDRs in both TSN2_NS and TSN2_HS interactomes using two predictor algorithms: PONDR-FIT and PONDR-VSL2 (Peng et al., 2005; Xue et al., 2010). The analysis revealed significant enrichment of both interactomes in IDR-containing proteins (Figure 3A). Based on their intrinsic disorder (ID) content, proteins were classified as highly ordered (disorder score < 0.25), moderately disordered (disorder score between 0.25 and 0.5) and highly disordered (disorder score > 0.5). According to PONDR-VSL2, while as much as 93% of the entire *Arabidopsis* proteome is represented by moderately and highly disordered proteins, this frequency was increased further in both TSN2_NS and TSN2_HS interactomes, reaching 99.4% and 100%, respectively (Figure 3B and Supplemental Figure S6).

**Figure 3.**
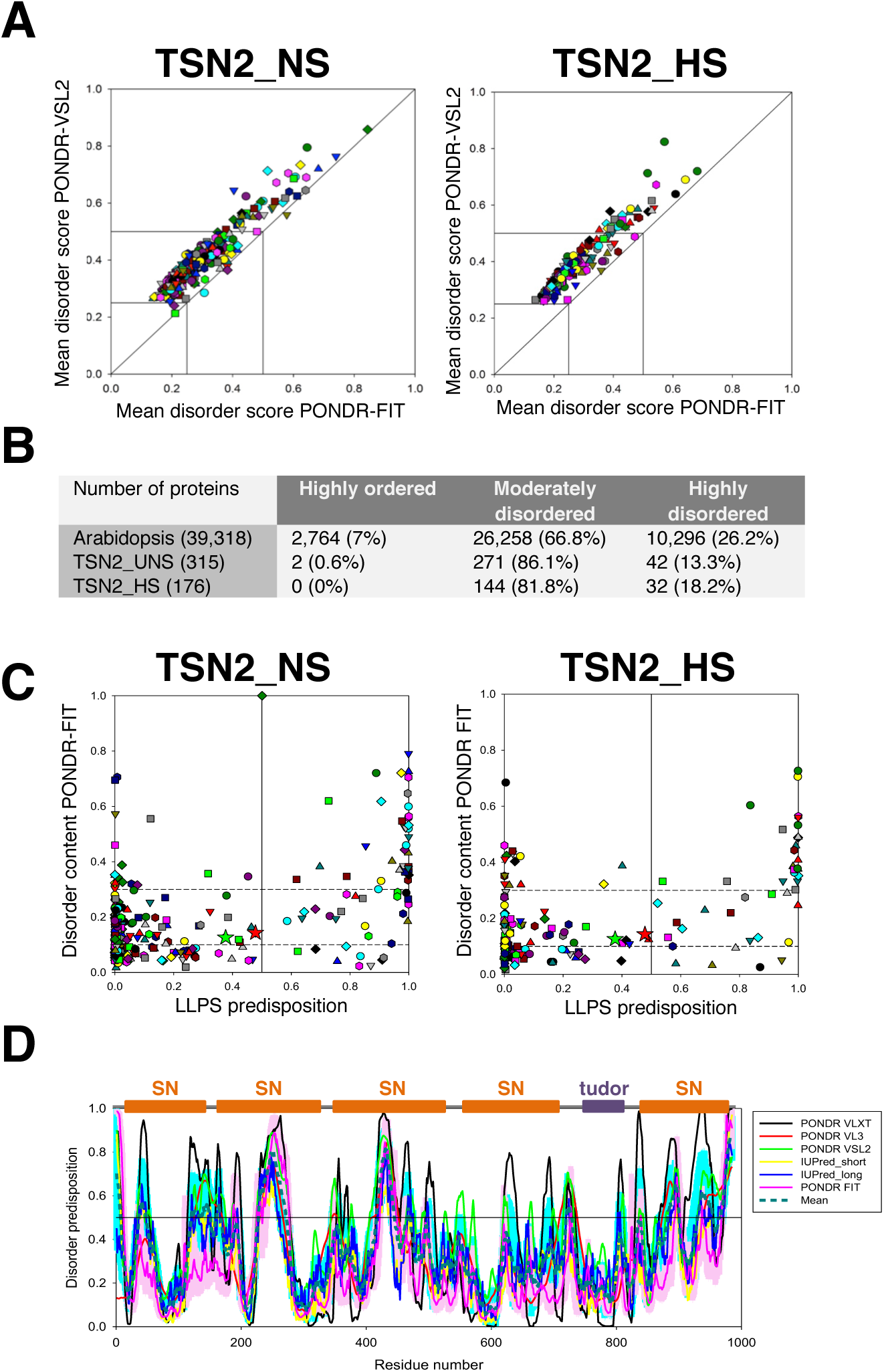
TSN2 interactomes are enriched in IDR-containing proteins. ***A***, Per-protein propensities for disorder (average of the corresponding per-residue propensities) evaluated by PONDR-FIT (*x*-axis) and by PONDR-VSL2 (y-axis) for TSN2_NS and TSN2_HS interactomes. ***B***, Classification of proteins from the whole *Arabidopsis* proteome and TSN2_NS and TSN2_HS interactomes based on their ID content. ***C***, Correlation between the ID content evaluated by PONDR-FIT (y-axis) and predisposition to undergo LLPS (*x*-axis). ***D***, Disorder profiles of TSN2 generated by PONDR-VLXT, PONDR-VL3, PONDR-VSL2, IUPred-short, IUPred-long and PONDR-FIT and a consensus disorder profile (based on mean values of six predictors). SN, staphylococcal nuclease domain.

Next, we evaluated the correlation between ID content and predisposition to undergo LLPS. As shown in Figure 3C ∼ 20% of proteins from both TSN2_NS (56/315) and TSN2_HS (36/177) bear propensity for LLPS (score > 0.5), and most of them are moderately (disorder score between 0.1 and 0.3) or highly (disorder score > 0.3) disordered. Taken together, the ubiquitous occurrence of IDRs among the TSN2-interacting proteins reinforces the view that TSN is a scaffolding factor seeding SG protein complexes.

In mammalian cells, highly disordered proteins such as TIA-1 are required to promote SG formation via LLPS (Protter and Parker, 2016; Rayman et al., 2018). Considering this fact as well as that TSN was shown to modulate the integrity of SGs in *Arabidopsis* (Gutierrez-Beltran et al., 2015), we evaluated the per-residue ID propensities of TSN2 using a set of six commonly used predictors, including PONDR-VLXT, PONDR-VL3, PONDR-VSL2, IUpred_short, IUpred_long and PONDR-FIT (Meng et al., 2015). Figure 3D shows that TSN2 is expected to have several (11 if averaged for six predictors) disordered regions (score above 0.5). Thus, the SN domains of TNS2 are predicted to be highly disordered, whereas the tudor region is predicted to be one of the most ordered parts of this protein. This observation was confirmed using D^2^P^2^ database providing information about the predicted disorder and selected disorder-related functions (Supplemental Figure S7A)(Oates et al., 2013). Notably, similar results were obtained for TSN1 protein isoform that is considered to be functionally redundant with TSN2 (Supplemental Figure S7) (dit Frey et al., 2010). Taken together, above results prompt us to believe that TSN proteins may recruit SG components via IDR regions, promoting in this way rapid coalescence of microscopically visible SGs upon stress exposure.

### TSN-interacting proteins co-localize with TSN2 in cytoplasmic foci

To ascertain the SG localization of TSN2-interacting proteins identified by mass spectroscopy, we shortlisted the 16 most interesting proteins from the HS-independent class of interactors (Figure 2C and 4A). The short list included homologues of key components of yeast and animal SGs (IF4E5, PAB4 and a 40S ribosomal subunit) and hypothetical plant-specific SG components with a key role in fundamental eukaryotic pathways (e.g. SnRK1 proteins, RH12, SKP1, MC1 and TCTP). First, we performed a co-localization study to investigate whether the shortlisted TSN-interacting proteins were translocated to TSN2 foci under stress. To this end, protoplasts were isolated from *Nicotiana benthamiana* (*N. benthamiana)* leaves co-transformed with RFP-TSN2 and individual GFP-TSN-interacting proteins. Co-transformation of the cytoplasmic protein GFP-ADH2 and the SG marker GFP-Ubp1c with RFP-TSN2 were used as a negative and positive control, respectively (Figure 4B, C). To quantify colocalization results, we calculated the linear Pearson (r_p_) and the nonlinear Spearman’s rank (r_s_) correlation coefficient (PSC) for the pixels representing the fluorescence signals in both channels (Figure 4C). The levels of co-localization can range from +1 for positive correlation to −1 for negative correlation (French et al., 2008). As shown in Figures 4B, C and S8, all shortlisted proteins co-localized with TSN2 in punctate foci under HS.

**Figure 4.**
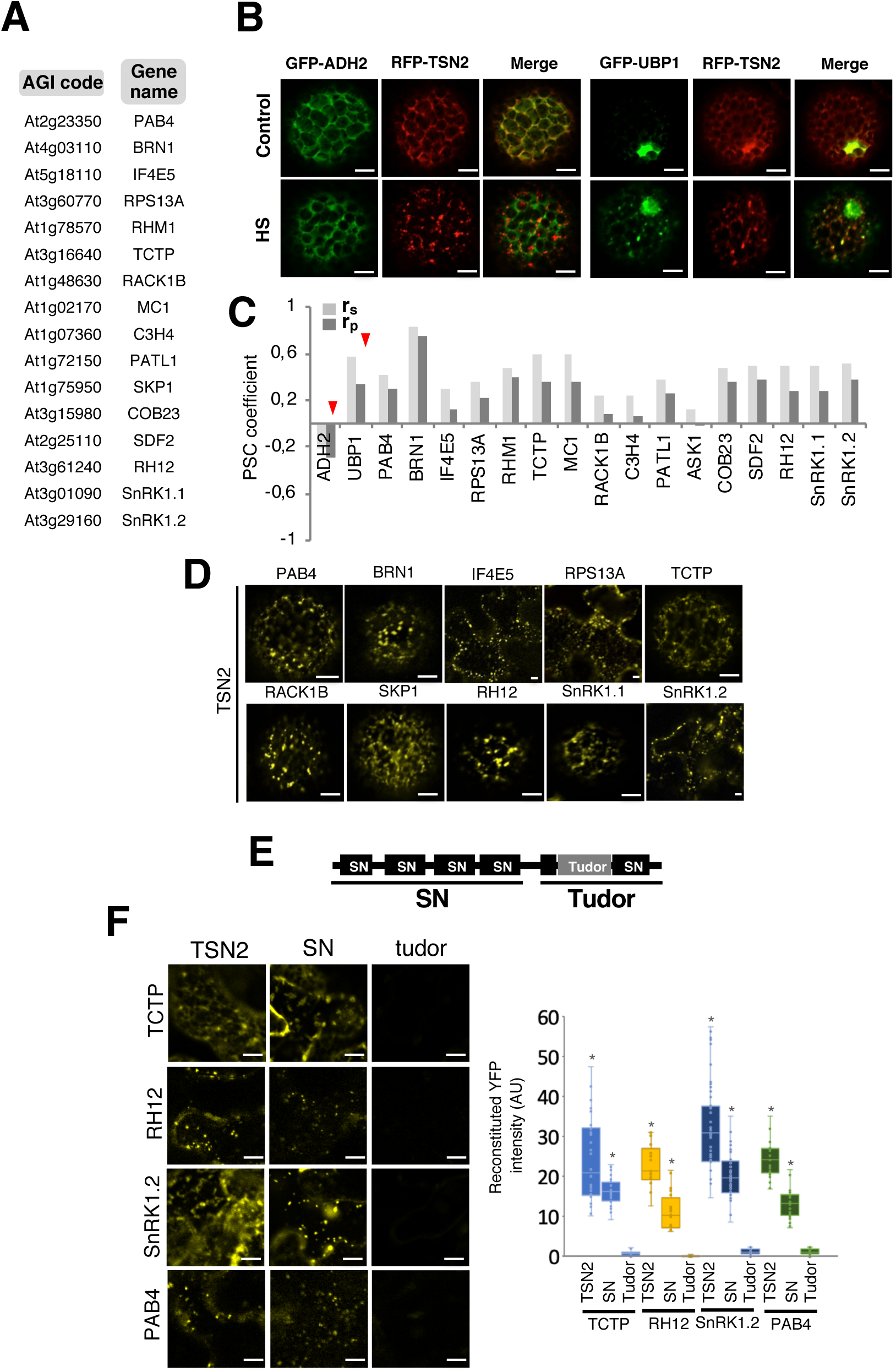
TSN2 and its interactors are localized in heat-induced SGs. ***A***, List of TSN-interacting proteins included in the co-localization analysis. ***B***, Co-localization of RFP-TSN2 (red) with GFP-ADH2 (negative control) and GFP-UBP1 (positive control) in *N. benthamiana* protoplasts incubated under control conditions (23°C) or at 39°C for 30 min (HS). Scale bars = 5 µm. ***C***, Pearson and Spearman coefficients (r_p_ and r_s_, respectively) of co-localization (PSC) of RFP-TSN2 with individual GFP-tagged TSN-interacting proteins listed in ***A*** and with both negative and positive control proteins (denoted by red arrowheads) under HS. ***D***, BiFC between cYFP-TSN2 and nYFP-TSN-interacting proteins in *N. benthamiana* protoplasts after HS (39°C for 30 min). Scale bars = 5 µm. ***E***, Schematic diagram of TSN protein domain organization depicting SN and Tudor regions. ***F***, BiFC between cYFP-TSN2 (full-length), cYFP-SN or cYFP-Tudor and nYFP-TSN-interacting proteins in *N. benthamiana* protoplasts after HS (39°C for 30 min). Scale bars = 10 µm. Chart shows quantification of the reconstituted YFP signal. AU, arbitrary units. Data are means ± SE of BiFC signal level measured in three independent experiments each containing seven individual measurements. Asterisks indicate significantly different fluorescence intensity compared with the corresponding Tudor (p<0.05, ANOVA).

Next, to elucidate whether these proteins interact with TSN2 in the heat-induced SGs, we performed bimolecular fluorescence complementation (BiFC) analysis with 10 of 16 shortlisted proteins. This analysis confirmed that all proteins indeed interacted with TSN2 in cytoplasmic foci under stress (Figure 4D). Notably, we observed that, in line with our proteomics data, TSN2 interacted with its partners also under normal (NS) conditions (Supplemental Figure S9). Taken together, these findings suggest that a subset of proteins that respond to stress by re-localization to punctate foci interact with TSN and are thus new candidates for SG-associated components in plants.

### N-terminally situated SN domains of TSN participate in the interaction with SG proteins

To investigate whether the interaction of TSN with SG proteins presents any domain preference, we performed a BiFC assay with full-length TSN2 or either SN region (tandem repeat of four N-terminally located SN domains) or Tudor (Tudor and the fifth SN domains) region (Figure 4E) fused to cYFP and four different SG-associated TSN-interacting proteins fused to nYFP in heat-stressed *N. benthamiana* leaves (Figure 4F). The results have revealed reconstitution of fluorescent signal in the experiments with all four TSN interactors in case of both full-length TSN2 and SN region, whereas none of the TSN interactors could form a complex with Tudor region (Figure 4F). Furthermore, expression of either full-length TSN2 or SN region yielded identical, punctate BiFC localization pattern. We conclude that tandem repeat of four SN domains confers TSN ability to recruit partner proteins to SGs.

### Arabidopsis SG components are common targets of TSN1 and TSN2 isoforms

*Arabidopsis* TSN1 and TSN2 proteins have been previously considered as functionally redundant (dit Frey et al., 2010; Gutierrez-Beltran et al., 2015). To investigate whether this redundancy is conserved at SG level, we isolated TSN1 interactome from unstressed plants using the same TAPa procedure as described above for TSN2 (Supplemental Figure S1B, S2, S3B). As a result, we obtained TSN1_NS pool comprised of 270 protein hits (Supplemental Figure S4 and Supplemental Table 1). Out of these, 108 (40%) were TSN1-specific, whereas the remaining, larger fraction (164 proteins, 60%) represented common interactors of TSN1 and TSN2, reflecting their functional redundancy (Figure 5A). Notably, the pool of common interactors of TSN1 and TSN2 was enriched in SG proteins, including core, evolutionarily conserved components such as PAB4, 40S ribosomal subunits, DEAD-box helicases or CCT proteins (group 1, Figure 5A). In addition to known SG homologs in either human or yeast, the TSN1/2 pool contains proteins found in the previously characterized *Arabidopsis* Rbp47b proteome (group 2), as well as novel plant SG components verified through either co-localization or BiFC with TSN2 or both methods (group 3, Figure 4A, C, D; Supplementary Figure S8).

**Figure 5.**
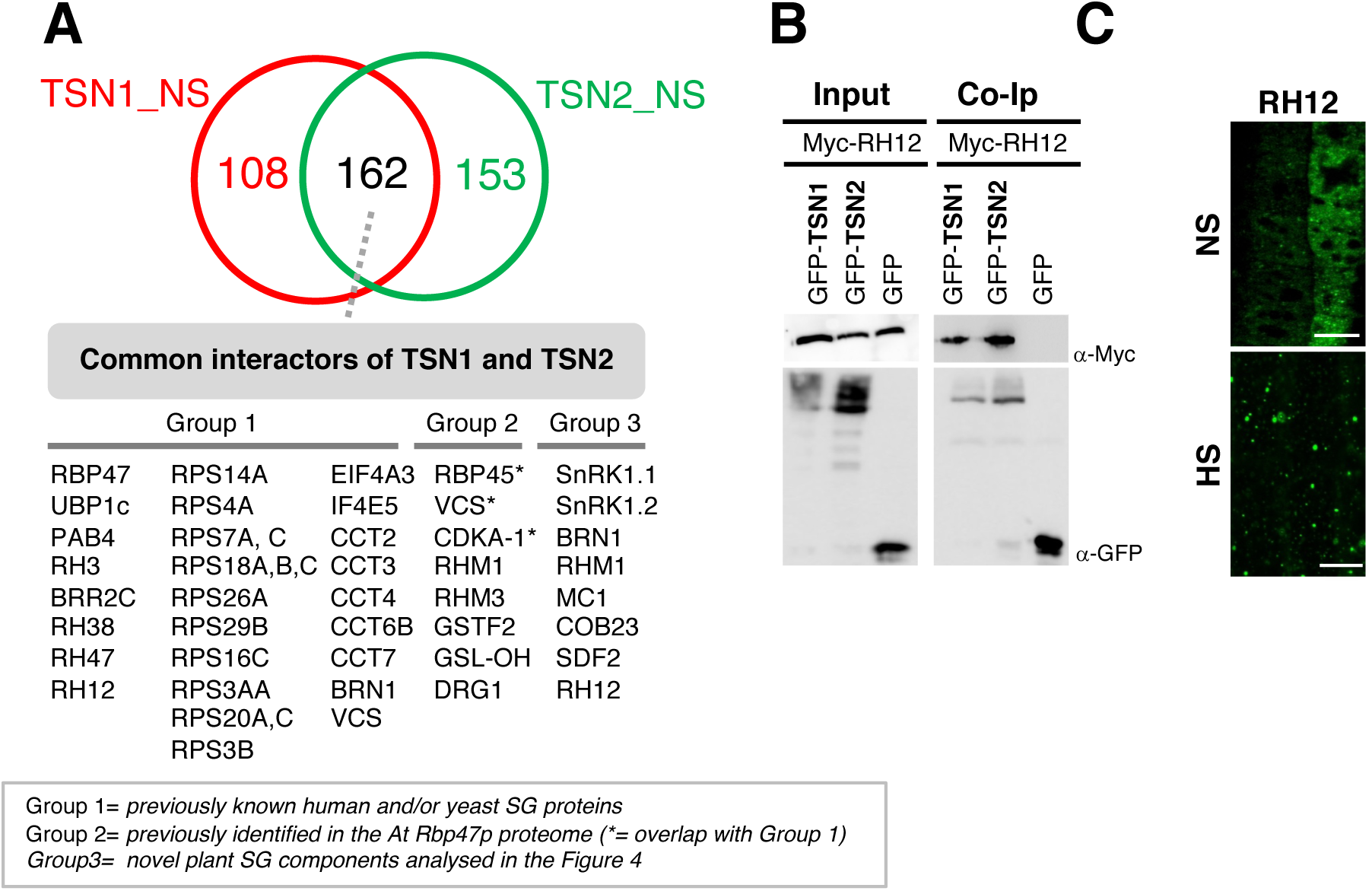
Proteomes of *Arabidopsis* TSN1 and TSN2 largely overlap. ***A,*** Venn diagram showing overlap between TSN1 and TSN2 interactomes isolated by TAPa from *Arabidopsis* plants grown under no stress (NS) conditions. Common interactors of TSN1 and TSN2 are classified into three groups. ***B,*** Co-Ip of TSN isoforms and RH12 in protein extracts prepared from *N. benthamiana* leaves agro-infiltrated with GFP-TSN1 or GFP-TSN2 and Myc-RH12. GFP was used as a negative control. Co-Ip was analysed by immunoblotting using α-Myc and α-GFP. *C*, Localization of RH12 in root cells of 5-old-day *Arabidopsis* seedlings expressing GFP-RH12 under control of the native promoter. The seedlings were grown under 23°C (NS) or incubated at 39°C for 40 min (HS). Scale bars = 10 µm.

To corroborate proteomics results, we have chosen DEAD-box ATP-dependent RNA helicase 12 (RH12), as a common interactor of TSN1 and TSN2. First, we confirmed molecular interaction between two isoforms of TSN and RH12 by coimmunoprecipitation (Co-IP) in cell extracts from agro-infiltrated *N. benthamiana* leaves. As shown in Figure 5B, RH12 coimmunoprecipitated with both TSN1 and TSN2 but did not with GFP, used as a negative control in this experiment. Second, we produced *Arabidopsis* lines stably expressing GFP-RH12 under native promoter and observed relocalization of the fusion protein to cytoplasmic foci under HS conditions in root tip cells (Figure 5C). Taken together, these data indicate that TSN1 and TSN2 are likely redundant in providing a scaffold platform for the recruitment of a wide range of SG components in *Arabidopsis* plants.

### Isolation of salt stress-induced TSN2 interactome and candidate core SG components

Previously it was shown that TSN2 is re-localized to SGs under salt stress (Yan et al., 2014). To investigate how TSN2 interactome is affected by salt stress, we isolated TSN2_NaCl interactome from salt-stressed *Arabidopsis* plants using our standard TAPa purification procedure. The resulting TSN2_NaCl pool included significantly lower number of protein hits (44 hits), as compared to both TSN2_NS and TSN2_HS pools hits (Figure 6A; Supplemental Figure S4 and Supplemental Table 1). Fifty two percent (23/44) of the protein hits were classified as NaCl stress-independent TSN2 interactors, as they also appeared in the TSN2-NS dataset, including many well-characterized SG proteins (e.g., Rbp47, UBP1, PAB4, several helicases and 40S ribosomal subunits). Interestingly, no known SG component were found among NaCl stress-dependent TSN2 interactors (Supplemental Table 1), suggesting a new role for TSN2 under salt stress that could be explored in future studies.

**Figure 6.**
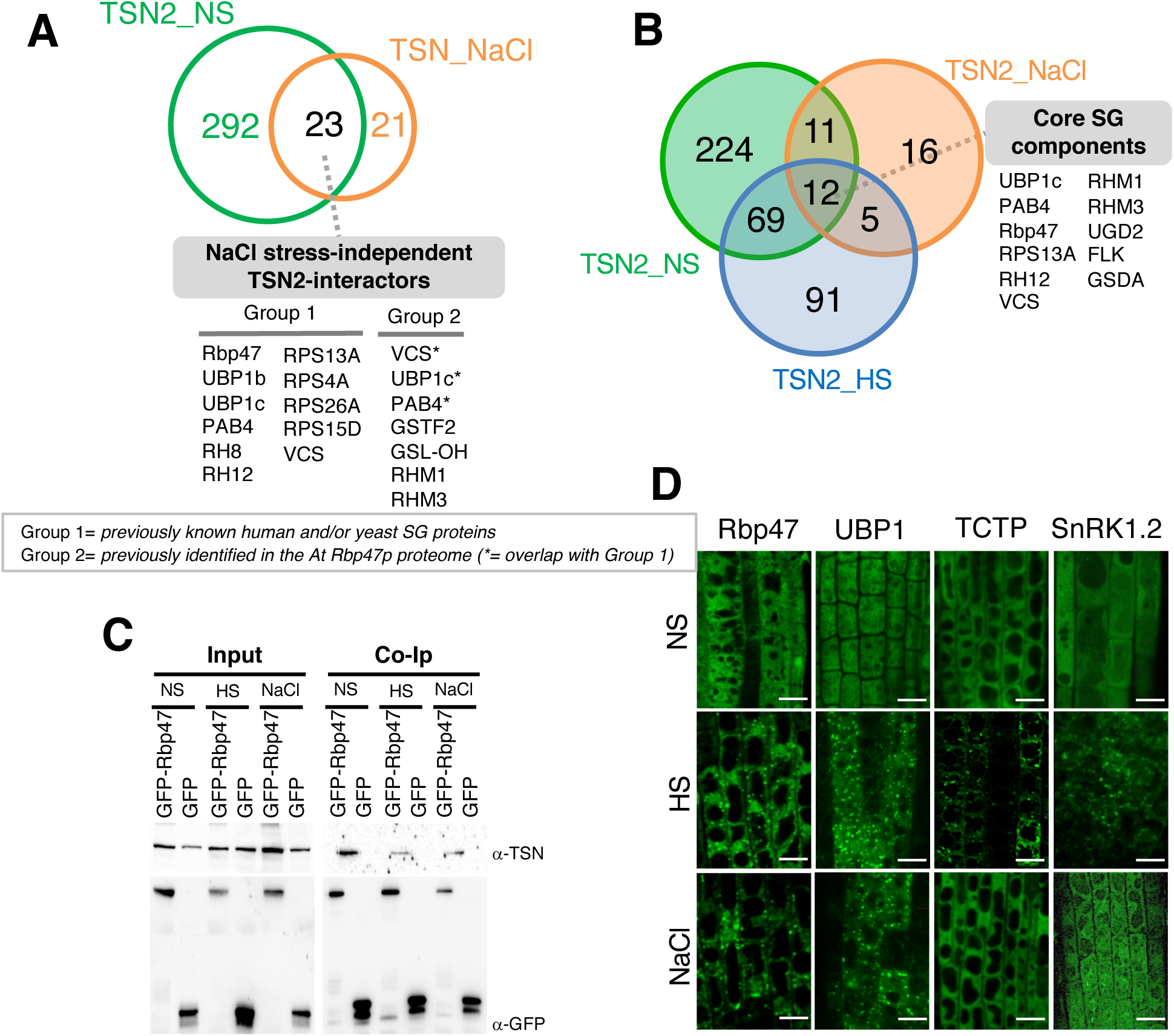
Characterization of presumed *Arabidopsis* SG core proteins. ***A***, Venn diagram showing comparison between TSN2_NS and TSN2_NaCl pools. Common TSN2 interactors are classified into two groups, based on their known localization to SGs in non-plant (Group 1) or plant (Group 2) organisms. ***B***, Venn diagram showing comparison between TSN2_NS, TSN2_HS and TSN2_NaCl pools. Eleven common TSN2 interactors are ascribed as presumed core SG components. ***C***, Co-Ip of TSN and Rbp47 in protein extracts prepared from *Arabidopsis* seedlings expressing Pro35S:GFP-Rbp47 and grown under no stress (C; NS), HS (39°C for 40 min) or salt stress (150 mM NaCl for 40 min) conditions. The GFP-expressing line was used as a negative control. Endogenous TSN (107 KDa) was detected in total fractions (Input) and fractions co-immunoprecipitated (Co-Ip) with Rbp47 but not with free GFP in all three conditions. Co-Ip was analysed by immunoblotting using α-TSN and α-GFP. ***D***, Localization of Pro35S:GFP-Rbp47, Pro35S:GFP-UBP1, Pro35S:GFP-TCTP and Pro35S:GFP-SnRK1.2 in root cells of 5-old-day *Arabidopsis* seedlings. The seedlings were grown under 23°C (NS) or incubated at 39°C for 40 min (HS) or with 150 mM NaCl for 40 min (NaCl). Scale bars = 10 µm

A broader comparative analysis including all three pools of TSN2 interactors, i.e. TSN2_NS, TSN2_HS and TSN2_NaCl yielded eleven proteins present in all three pools and presumably representing core SG components constitutively bound to the TSN platform (Figure 6B). To validate this assumption, we performed Co-IP using protein extracts prepared from 7-day-old *Arabidopsis* seedlings expressing GFP-Rbp47 (a putative core SG protein fused to GFP) and exposed to heat or salt stress (Rayman et al., 2018). As shown in Figure 6C, TSN co-immunoprecipated with GFP-Rbp47 under both types of stresses, as well as in the absence of stress. To further corroborate our result by *in planta* observations, we produced *Arabidopsis* lines stably expressing GFP-fused variants of Rbp47 and UBP1, two putative core SG proteins, and TCTP and SnRK1.2, two HS-dependent TSN2 interactors. Analysis of root tip cells revealed that while Rbp47b and UBP1 were localized to both HS- and NaCl-induced SG puncta, TCTP and SnRK1.2 exhibited punctate localization only under HS (Figure 6D).

### TSN and SGs confer heat-induced activation of SnRK1

We have found that SnRK1.1 and SnRK1.2, - two *Arabidopsis* homologues of the evolutionary conserved SNF1-related protein kinase 1, - are novel TSN-interacting proteins re-localized to SGs exclusively during HS (Figure 4 and Figure 6D). To dissect the functional relevance of TSN binding and SG localization of SnRK1.1 and SnRK1.2, we investigated whether HS and the presence of TSN could affect their kinase activity. To begin with, we corroborated the interaction with TSN2 using two different approaches. First, we performed co-immunoprecipitation of native TSN and GFP-SnRK1.2 in protein extracts prepared from heat-stressed *Arabidopsis* plants expressing GFP-SnRK1.2. We found that native TSN co-immunoprecipitated with GFP-SnRK1.2 but not with GFP, which was used as a negative control (Figure 7A). Second, a Förster resonance energy transfer (FRET) assay demonstrated that TSN2 directly interacts with SnRK1.2 in *N. benthamiana* leaves under HS (Figure 7B). Taken together, these findings confirm the *in vivo* TSN-SnRK1 interaction.

**Figure 7.**
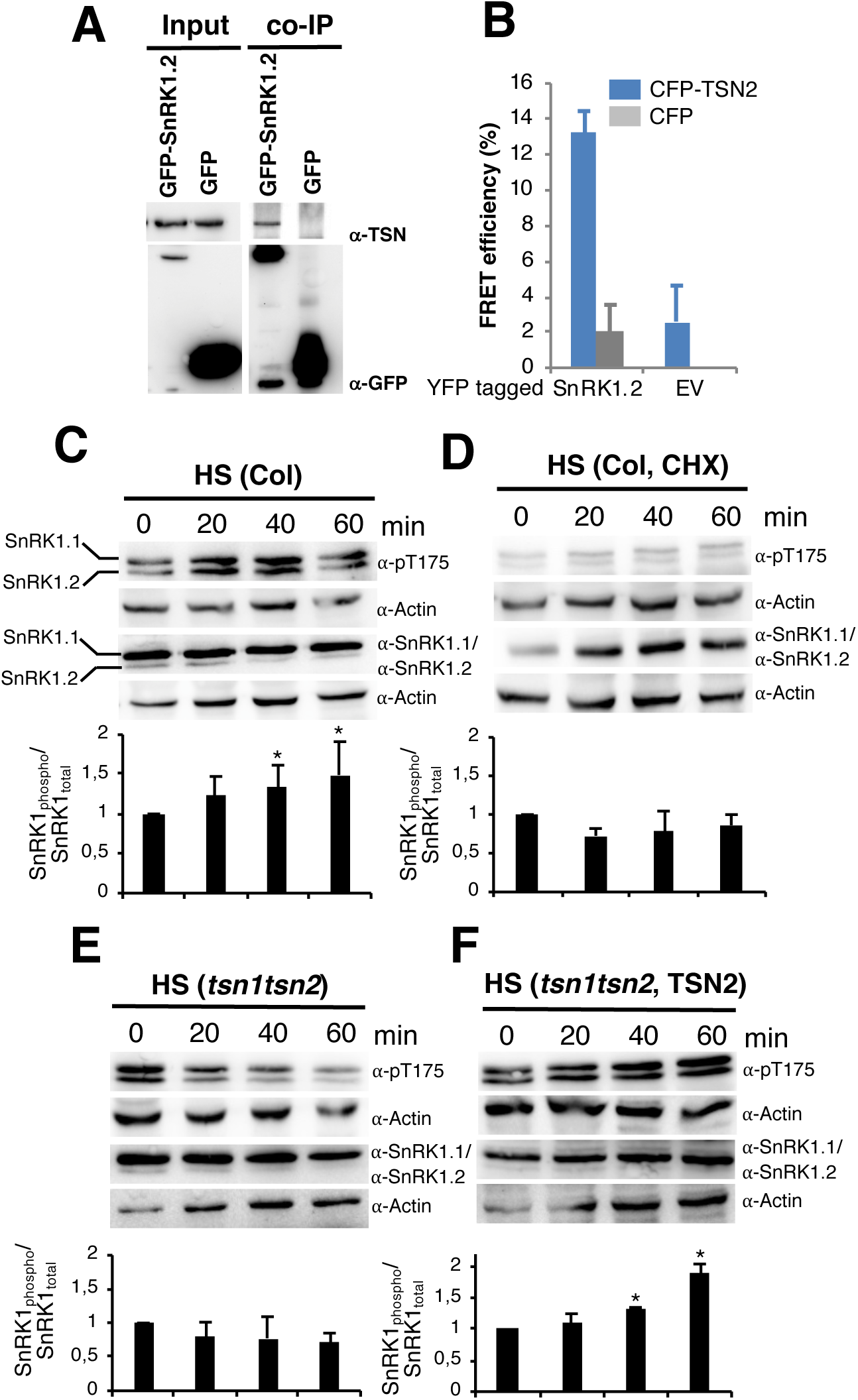
Activation of SnRK1 under HS depends on the presence of both TSN and SGs. ***A***, Co-Ip of TSN and SnRK1.2 in protein extracts prepared from *Arabidopsis* seedlings expressing Pro35S:GFP-SnRK1.2 and grown under heat stress condition (39°C for 40 min). The GFP-expressing line was used as a negative control. Endogenous TSN (107 KDa) was detected in the total fractions (Input) and in the fraction co-immunoprecipitated (Co-Ip) with SnRK1.2 but not with free GFP. Co-Ip was analysed by immunoblotting using α-TSN and α-GFP. ***B***, FRET assay of the indicated protein combinations using CFP-YFP pair in *N. benthamiana* leaves under HS (39°C for 40 min). EV, empty vector (negative control). Data show mean ± SD of 10 replicate measurements. The experiment was repeated three times with similar results. ***C-F***, Immunoblot analysis with α-SnRK1.1, α-SnRK1.2, α-p-T175 and α-Actin of protein extracts prepared from root tips of 10-day-old *Arabidopsis* Col **(*C, D*),** *tsn1tsn2 **(E)** or tsn1tsn2* expressing *ProTSN2:GFP-TSN2 **(F)*** seedlings heat-stressed (39°C) for 0, 20, 40 and 60 min. ***D***, The seedlings were pre-treated with cycloheximide (CHX, 200 ng/µL) and then subjected to HS. SnRK1 activity was determined as the ratio of phosphorylated to total SnRK1 protein. The data on charts in ***C-F*** show mean ratio of phosphorylated to total SnRK1 (both isoforms) integrated band intensity level ± SD from five experiments. Asterisks denote significant difference; Student’s t test, P < 0.05.

To determine whether SnRK1 activity is regulated *in vivo* by HS, we subjected 10-day-old Columbia (Col) *Arabidopsis* seedlings to 39°C for 0, 20, 40 and 60 min, and performed immunoblotting using α-phospho-AMPK Thr175 (α-pT175), which recognizes both SnRK1.1 (upper band 61.2 kDa) and SnRK1.2 (lower band 58.7 kDa) (Rodrigues et al., 2013; Nukarinen et al., 2016). As a control test, we confirmed the α-pT175 sensitivity using ABA treatment known to induce SnRK1 T175 phosphorylation (Figure S10) (Jossier et al., 2009). Time course analysis of the level of SnRK1 T175 phosphorylation under HS demonstrated that the two SnRK1 isoforms are rapidly activated by stress (Figure 7C). To exclude the possible bias due to fluctuations in the protein expression level, we conducted immunoblotting assays using α-SnRK1.1 and SnRK1.2, and found that the protein levels of both kinases were not affected by the treatment. To further correlate the SnRK1 activity with the formation of SGs, the seedlings were treated with cycloheximide (CHX), a drug blocking SG assembly in mammalian and plant cells (Gutierrez-Beltran et al., 2015; Wolozin and Apicco, 2015). As shown in Figure 7D, CHX treatment abrogated heat-induced phosphorylation of SnRK1 T175, suggesting that activation of SnRK1 isoforms is linked to the formation of heat-induced SGs.

To test whether TSN is involved in the regulation of the SnRK1 kinase activity, *tsn1tsn2* double mutant seedlings were exposure to 39°C for 0, 20, 40 and 60 min, and protein extracts were analyzed by immunoblotting with α-pT175. As in case of CHX treatment, TSN deficiency prevented heat-induced phosphorylation of SnRK1 T175 (Figure 7E). Accordingly, complementation of the *tsn1tsn2* double mutant with TSN2 under the native promoter resulted in full rescue of the heat-induced phosphorylation of SnRK1 T175 (Figure 7F), confirming that TSN is required for the activation of SnRK1 during HS.

## DISCUSSION

One of the earliest, evolutionary conserved events occurring upon stress perception and providing defence mechanism to promote cell survival is the assembly of SGs in the cytoplasm of eukaryotic cells (Thomas et al., 2011; Mahboubi and Stochaj, 2017). Molecular composition and regulation of SGs is a rapidly growing area, but most of the works done so far utilized animal or yeast models.

In a recent study, we found that TSN is stably associated with SGs in *Arabidopsis* (Gutierrez-Beltran et al., 2015). According to current hypothesis suggesting that the SG cores are relatively stable, while the SG shells are highly dynamic (Jain et al., 2016), we hypothesize that TSN is a SG core protein in plants. In the present study, we found that the *Arabidopsis* TSN proteins interact with numerous SG components and that most of these interactions take place under no stress condition (Figure 2). This finding, together with the fact that N-terminally situated SN domains are essential for these interaction, as well as SG-specific localization of TSN (Zhu et al., 2013; Gutierrez-Beltran et al., 2015), make it reasonable to propose the potential role of the SN domains as a docking platform maintaining a pre-existing state of SGs in plant cells (Figure 8).

**Figure 8.**
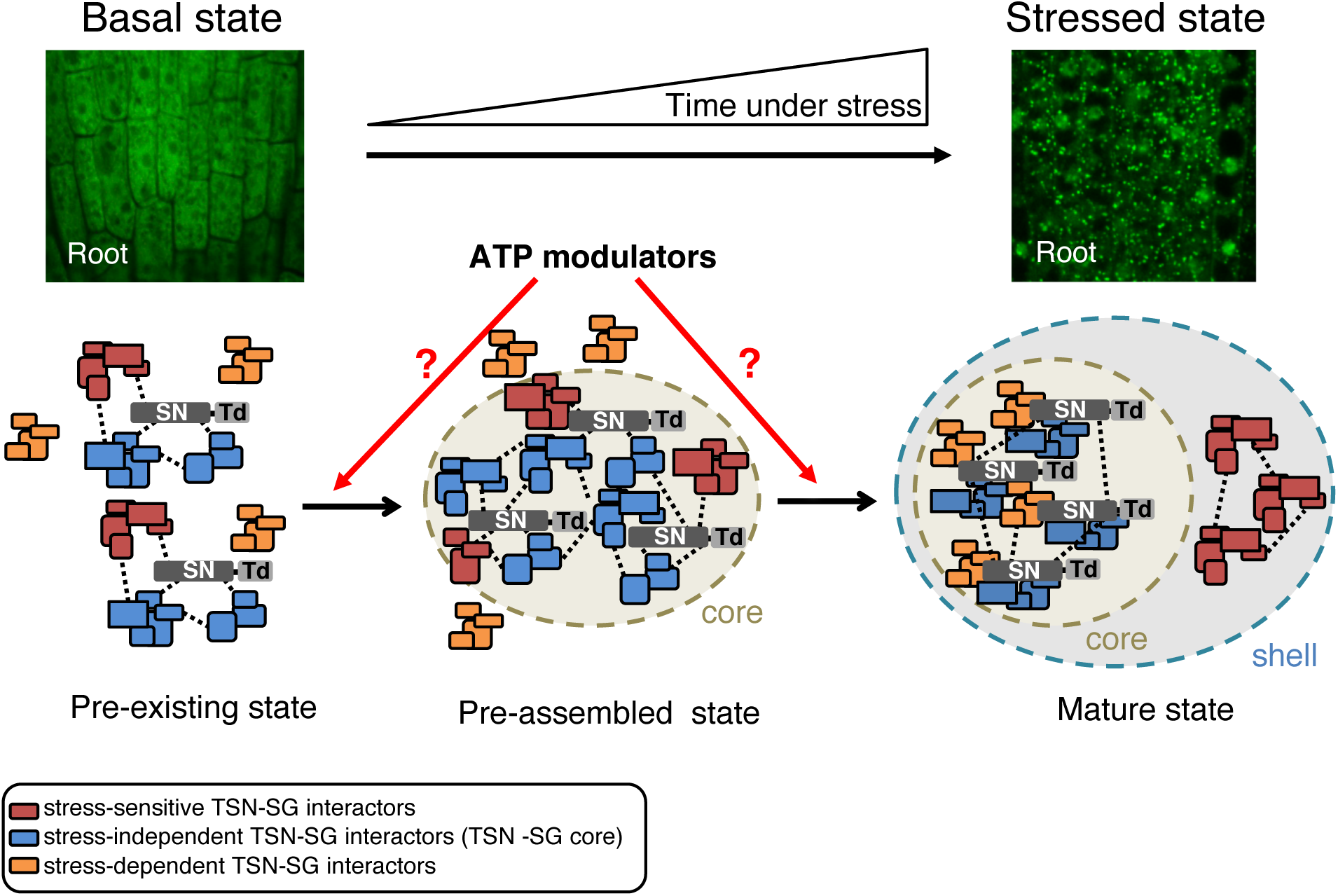
Schematic drawing showing maturation of SGs upon stress perception whereupon TSN serves a docking platform for SG components. Plant stress granules can by hypothesized to undergo two phases of assembly. The first phase of this model is formation of a pre-assembly state in which stress granules core is assembled by protein-protein interaction mediated by IDR from proteins present in a pre-existing TSN-SG complex. Second, formation of a larger microscopically visible stress granules (mature state) is constituted upon removing of stress-sensitive TSN interactors and incorporation of stress-dependent interactors. Dashed lines represent physical interactions between IDR-containing proteins. Red lines represent possible sites of activity of the ATP modulators. The upper part show 5-day-old Col seedlings expressing ProTSN2:TSN2-GFP grown under no stress (23°C, Basal state) or incubated for 40 min at 39°C (Stressed state).

There are several lines of evidence suggesting that assembly of mammalian and yeast SGs might be a highly regulated process controlled, at least in part, by ATP-dependent remodeling complexes (Protter and Parker, 2016). First, numerous energy-driven chaperones have been found in SG proteomes. Second, ATP is required for the formation of SGs (Jain et al., 2016). Therefore, ATP-dependent events mediated by ATPases, such as movement of mRNPs to sites of SG formation by motor proteins or remodeling of mRNPs to load required components could be imperative for promoting SG assembly. In this context, the interaction of chaperonin-containing T complex (CCT complex) with SG components has been suggested to be crucial for the proper assembly of SGs in yeast (Jain et al., 2016). Similarly, a mutation of DEAD-box helicase 1 (Ded1) that blocks its ATPase activity leads to retention of mRNAs in SGs and accumulation of these foci in the cytoplasm of yeast cells (Hilliker et al., 2011). In plants, the homologues of the yeast DEAD-box RNA helicase DHH1 is required for proper formation of stress granules (Chantarachot. et al., 2019). Considering that an enrichment of the TSN interactome in ATP-dependent remodeling complexes, including CCT proteins and DEAD-box RNA/DNA helicases occurs exclusively in the absence of stress stimulus (Figure 2C), we hypothesize that interaction between these proteins and TSN is necessary for the early steps of SG assembly in plants (Figure 8). Once the stress stimulus is perceived, the ATP-dependent remodeling complexes might detach from the TSN platform and aid in SG shell assembly.

The composition of SG proteome in animal and yeast cells is a highly variable characteristics influenced by a type of stress or cell type (Markmiller et al., 2018). However, certain proteins, e.g. G3BP1 and PAB, are constant SG constituents (Mahboubi and Stochaj, 2017). We estimated that up to 50% of TSN-interacting proteins may be recruited to *Arabidopsis* SGs in stress type-specific manner. In addition to a large resource of nearly 400 previously unknown plant candidate SG proteins for further validation, our study provides also a subset of proteins constantly interacting with TSN, regardless of stress or a type of stress. These proteins, including UBP1, PAB4, Rbp47 or RH12 can therefore be considered as core constituents of plant SGs (Figure 6B).

In non-plant species, one of the most enriched categories of molecular components of SGs are RBPs, regulating RNA transport, silencing, translation and degradation (Wolozin and Apicco, 2015). Likewise, RBPs accounted for 53% and 54% of TSN2_HS and TSN2_NS interactomes (Figure 2B), respectively, providing a further mechanistic explanation for the previously established role of TSN in mRNA stabilization and degradation (Gutierrez-Beltran et al., 2015). Yet, we found a high occurrence of ribosomal proteins, including components of 40S subunits, consistently with several studies showing a close association between ribosomes and SGs (Yang et al., 2014; Cary et al., 2015). Curiously, we have not found any PB-specific proteins, such as DCP or XRN, among *Arabidopsis* TSN interactors. In contrast, proteins such as VCS, PATL1 and several helicases, which have been localized to both SG and PB cytoplasmic foci (Youn et al., 2018) were identified in the TSN interactome, suggesting that similarly to yeast and animal models, SGs and PBs in plants share a part of their components.

A current, predominating model for SG assembly rests on LLPS driven by dynamic and promiscuous interactions among IDRs (Molliex et al., 2015; Rayman et al., 2018). In this context, the overexpression of the prion-like domain (a type of IDR) of the mammalian SG core protein TIA-1 was sufficient to promote the formation of SGs (Kedersha et al., 1999; Gilks et al., 2004). In *Arabidopsis* plants, the prion-like domains of the TIA-1 homologues UBP1 and RBP47 were required for protein targeting to SGs (Weber et al., 2008). Our data further demonstrate that almost all *Arabidopsis* TSN-interacting proteins are disordered (Figure 3B), and ∼ 20% of them are predisposed for LLPS (Figure 3C). Lastly, TSN protein itself is highly disordered, with the most IDRs located within its five SN domains (Figure 3D), four of which, situated at the N-terminus were shown to confer TSN interaction with partner proteins, SG localization and cytoprotective function in both mammalian and plant cells (Figure 4F; (Gao et al., 2015; Gutierrez-Beltran et al., 2015). Taken together, these results have two important implications. First, the function of IDRs in SG condensation is conserved in plants. Second, the IDRs of TSN-dependent protein complexes may underpin SG functions.

It is well known that numerous stress- and nutrient-signaling pathways converge on SGs (Kedersha et al., 2013; Mahboubi and Stochaj, 2017). Our study has established two homologues of the evolutionary conserved signaling protein SnRK1 (SnRK1.1 and SnRK1.2; also known as KIN11 and KIN10, respectively) as TSN interactors. SnRK1 (AMPK in mammals and Snf1 in yeast) has been extensively studied as one of the key regulators of the target of rapamycin (TOR) (Shaw, 2009). In plants, SnRK1 and TOR proteins play central and antagonistic roles as integrators of transcription networks in stress and energy signaling (Baena-Gonzalez et al., 2007). Thus, while SnRK1 signaling is activated during stress and energy limitation, TOR promotes growth and biosynthetic processes in response to energy availability (Baena-Gonzalez and Hanson, 2017; Carroll and Dunlop, 2017). Although it has been demonstrated that the mammalian orthologue (AMPK) is a *bona fide* SG component involved in the regulation of SG biogenesis (Mahboubi et al., 2015), there is no evidence connecting SnRK1 activation and SGs. Here we demonstrate that formation of SGs and the presence of TSN are both necessary for SnRK1 activation in response to HS (Figure 7). It has been shown that mammalian mTOR is translocated to SGs under stress, leading to its inactivation (Heberle et al., 2015). While there is no evidence so far that TOR is a component of SGs in plants, we detected a TOR downstream effector RPS6 among TSN-interacting proteins (Figure 2C, Supplemental Table 1). We thus speculate that SGs and their integral constituent protein TSN might play a crucial role in the regulation of the SnRK1-TOR module; however further work is required to decipher mechanistic details and physiological roles of this regulation.

Based on our current results, we propose a three-step working model for the TSN-dependent biogenesis of plant SGs that encompasses formation of the pre-existing TSN-SG complex, as the first, primordial step crucial for proper SG assembly (Figure 8). A few minutes after stress exposure, a high-density protein-protein interaction network mediated by IDRs of stress-independent TSN interactors induces LLPS, leading to the formation of a pre-assembled state. Since TSN is a core component (Gutierrez-Beltran et al., 2015), one possibility is that formation of stress granule core takes part first, followed by the assembly of a shell around this core in which detachment of stress-sensitive TSN interactors and incorporation of stress-dependent interactors accomplishing SG maturation process. ATP modulators present in the stress-sensitive pool such as CCT or DEAD-box RNA/DNA helicases, could be critically required for both transition steps.

## EXPERIMENTAL PROCEDURES

### Plant Material and Molecular Biology

The T-DNA *tsn1tsn2* double mutant for *TSN1* and *TSN2*, in the Landsberg erecta (Ler) and Columbia (Col) backgrounds, respectively, was isolated as shown previously (Gutierrez-Beltran et al., 2015). The mutant was five times back-crossed with wild-type (WT) Col plants to generate an isogenic pair. Finally, both *tsn1tsn2* mutant and WT plants were selected from F5. All oligonucleotide primers used in this study are shown in Supplemental Table 3. *TSN1* (no stop codon egb1/egb29; with stop codon egb1/egb28), *TSN2* (no stop codon egb4/egb6; with stop codon egb4/egb5), and *GFP* (egb19/egb20) were amplified by PCR from pGWB6 and resulting cDNA sequences were introduced in pDONR/Zeo using Gateway technology (Invitrogen). For expression of C-terminal TAPa fusion under the control of (2x) 35S promoter, *TSN1*, *TSN2* and *GFP* cDNAs were introduced into the destination vector pC-TAPa (Rubio et al., 2005). cDNA clones of TSN-interacting proteins in the Gateway compatible vector pENTR223 were obtained from the ABRC stock center (Yamada et al., 2003). For expression of N-terminal GFP and RFP fusions under the control of 35S promoter, cDNAs encoding TSN2 and TSN-interacting proteins were introduced into the destination vectors pMDC43 and pGWB655, respectively (Curtis and Grossniklaus, 2003). For BiFC assay, cDNAs for TSN2, TSN-interacting proteins, as well as SN and Tudor regions were cloned into pSITE-BiFC destination vectors (Martin et al., 2009). For FRET experiments, cDNAs for TSN2 and SnRK1.2 were introduced into pGWB642 (YFP) and pGWB645 (CFP) destination vectors (Nakamura et al., 2010). All plasmids and derived constructs were verified by sequencing using the M13 forward and reverse primers.

### Tandem Affinity Purification

Fully expanded leaves from *Arabidopsis* Col transgenic plants expressing TSN-TAPa and GFP-TAPa and grown in the greenhouse for 18 days in 18:6 light/dark conditions at 22 °C (NS), 39°C for 40 min (HS) and 200 mM NaCl_2_ for 5 h (NaCl) were harvested (15 g, fresh weight) and ground in liquid N_2_ in 2 volumes of extraction buffer (50 mM Tris–HCl pH 7.5, 150 mM NaCl, 10% glycerol, 0.1% Nonidet P-40 and 1x protease inhibitor cocktail; Sigma-Aldrich) and centrifuged for 12,000 *g* for 10 min at 4°C. Supernatants were collected and filtered through two layers of Miracloth (Calbiochem). Plant extracts were incubated with 700 µL IgG beads (Amersham Biosciences) for 4-5 h at 4°C with gentle rotation. After centrifugation at 250 *g* for 3 min at 4°C, the IgG beads were recovered and washed three times with 10 mL of washing buffer (50 mM Tris-HCl, pH 7.5, 150 mM NaCl, 10% glycerol, and 0.1% Nonidet P-40) and once with 5 mL of cleavage buffer (50 mM Tris-HCl, pH 7.5, 150 mM NaCl, 10% glycerol, Nonidet P-40, and 1 mM DTT). Elution from the IgG beads was performed by incubation with 15 µL (40 units) of PreScission protease (Amersham Biosciences) in 5 mL of cleavage buffer at 4°C with gentle rotation. Supernatants were recovered after centrifugation at 250 *g* for 3 min at 4°C and stored at 4°C. The IgG beads were washed with 5 mL of washing buffer, centrifuged again, and the eluates pooled. Pooled eluates were transferred together with 1.2 mL of Ni-NTA resin (Qiagen, Valencia, CA, USA) into a 15 mL Falcon tube and incubated for 2 h at 4°C with gentle rotation. After centrifugation at 250 *g* for 3 min at 4°C, the Ni-NTA resin was washed three times with 10 mL washing buffer. Finally, elution was performed using 4 mL of imidazole-containing buffer (50 mM Tris–HCl pH 7.5, 150 mM NaCl, 10% glycerol, 0.1% Nonidet P-40, 200 mM imidazole). All the steps in the purification procedure were carried out at 4°C. For each large-scale TAPa purification, three TAPa plant samples (15 g, fresh weight each) were processed in parallel as described above. Final eluates were pooled together, proteins were precipitated using TCA/Acetone extraction and 100 µg of protein was digested according to the FASP method (Wisniewski et al., 2009). Three and two biological replicates were performed for isolating TSN interactomes from unstressed and stressed plants, respectively.

### Liquid chromatography and mass spectrometry analysis

Peptides were analyzed using EASYnano-LC 1000 on a Q Exactive Plus Orbitrap mass spectrometer (Thermo Scientific). Peptides were separated on a pre-column 75 µm × 2 cm, nanoViper, C18, 3 µm, 100 Å (Acclaim PepMap 100) and analytical column 50 µm × 15 cm, nanoViper, C18, 2 µm, 100 Å (Acclaim PepMap RSLC) at a flow rate of 200 nL/min. Water and ACN, both containing 0.1% formic acid, were used as solvents A and B, respectively. The gradient was started and kept at 0-35% B for 0-220 min, ramped to 35-45% B over 10 min, and kept at 45-90% B for another 10 min. Operate the mass spectrometer in the data-dependent mode (DDA), to automatically switch between full scan MS and MS/MS acquisition. Acquire survey full scan MS spectra from 200 to 1800 *m/z* in the Orbitrap with a resolution of R = 70,000 at *m/z* 100. For data dependent analysis, the top 10 most abundant ions were analyzed by MS/MS, while +1 ions were excluded, with a normalized collision energy of 32%. The raw data were searched with the Sequest HT node of Proteome Discoverer 1.4. The Uniprot database (*Arabidopsis* TAIR10), was utilized for the searches. All protein identification results were filtered to include only high confidence peptides with peptide mass deviation 2 and a minimum of 2 unique peptides per protein and score thresholds to attain an estimated false discovery rate of **∼**1% using a reverse decoy database strategy (Chittum et al., 1998).

### Plant and Protoplast Transformation

*Arabidopsis* Columbia (Col) plants were transformed as described previously (Clough and Bent, 1998) using *Agrobacterium tumefaciens* (Agrobacterium) strain GV3101. In Figure 5 and 6, plants from the T2 and T3 generations were used. Transgenic plants were confirmed by genotyping. For transient expression in *N. benthamiana* mesophyll cells, Agrobacterium strain GV3101 was transformed with the appropriate binary vectors by electroporation as described previously (Gutierrez-Beltran et al., 2017). Agrobacterium-positive clones were grown in Luria-Bertani until reaching OD_600_= 0.4 and were pelleted after centrifugation at 3,000 *g* for 10 min. Cells were resuspended in MM (10 mM MES, pH 5.7, 10 mM MgCl_2_ supplemented with 0.2 mM acetosyringone) until OD_600_= 0.4, incubated at room temperature for 2 h, and infiltrated in *N. benthamiana* leaves using a 1 mL hypodermic syringe. Leaves were analyzed after 48 h using a Zeiss 780 confocal microscope with the 40x objective. The excitation/emission wavelength was 480/508 nm for GFP, and 561/610 nm for RFP.

Protoplasts were isolated from leaves of 15- to 20-day-old *N. benthamiana*, transiently expressing the corresponding fluorescent proteins, as described previously (Wu et al., 2009). The cell walls were digested in enzymatic solution containing 1% (w/v) Cellulose R-10, 0.25% (w/v) Macerozyme R-10, 20 mM MES-HOK pH 5.7, 400 mM Mannitol, 10 mM CaCl_2_, 20 mM KCl, 0.1 % (w/v) Bovine serum albumin (BSA) for 60 min. Protoplasts were separated from debris by centrifugation (100 *g*, 3 min, 4°C), washed two times with ice-cold W5 buffer (154 mM NaCl, 125 mM CaCl_2_, 5 mM KCl and 2 mM MES-KOH pH 5.7) and resuspended in ice-cold W5 buffer at a density of 2.5 × 10^5^ protoplasts mL^−1^. The protoplast suspension was incubated for 15 min on ice before heat stress.

### Bimolecular Fluorescence Complementation (BiFC)

For BiFC assays, Agrobacterium strains GV3101 carrying *cYFP-TSN2 cYFP-SN or cYFP-Tudor* and the corresponding *nYFP-TSN-interacting proteins* were co-infiltrated into *N. benthamiana* leaves (OD_600_=0.3). Fluorescence images were obtained 48 h after infiltration using a Leica TCS Sp2/DMRE confocal microscope, with excitation wavelength 514 nm. Transient expression of proteins in *N. benthamiana* leaves via agroinfiltration was performed as previously described (Gutierrez-Beltran et al., 2017).

### Immunocytochemistry and Imaging

Five-day-old *Arabidopsis* roots were fixed for 60 min at room temperature with 4% (w/v) paraformaldehyde in 50 mM PIPES, pH 6.8, 5 mM EGTA, 2 mM MgCl_2_, and 0.4% Triton X-100. The fixative was washed away with phosphate buffered saline buffer supplemented with Tween 20 (PBST) and cells were treated for 8 min at room temperature with a solution of 2% (w/v) Driselase (Sigma-Aldrich) in 0.4 M mannitol, 5 mM EGTA, 15 mM MES pH 5.0, 1 mM PMSF, 10 µg mL^−1^ leupeptin, and 10 µg mL^−1^ pepstatin A. Thereafter, roots were washed twice, 10 min each, in PBST and then in 1% (w/v) BSA in PBST for 30 min before overnight incubation with a primary antibody (rabbit α-Myc diluted 1:500). The specimens were then washed three times for 90 min in PBST and incubated overnight with goat anti-rabbit fluorescein isothiocyanate (FITC) conjugated secondary antibody diluted 1:200. After washing in PBST, the specimens were mounted in Vectashield mounting medium (Vector Laboratories).

### Förster Resonance Energy Transfer (FRET)

FRET was performed using Zeiss 780 laser scanning confocal microscope and a plan-apochromat 20x/0.8 M27 objective. FRET acceptor photobleaching mode of Zeiss 780 ZEN software was used, with the following parameters: acquisition of 10 pre-bleach images, one bleach scan, and 80 post-bleach scans. Bleaching was performed using 488, 514 and 561 nm laser lines at 100% transmittance and 40 iterations. Pre- and post-bleach scans were at minimum possible laser power (0.8 % transmittance) for the 458 nm or 514 nm (4.7%) and 5% for 561 nm; 512 × 512 8-bit pixel format; pinhole of 181 µm and zoom factor of 2.0. Fluorescence intensity was measured in the ROIs corresponding to the bleached region. One ROI was measured outside the bleached region to serve as the background. The background values were subtracted from the fluorescence recovery values, and the resulting values were normalized by the first post-bleach time point. Three pre-bleach and three post-bleach intensities were averaged and used for calculations using the formula *FRET*_eff_ = (D_post_-D_pre_)/D_post_, where D is intensity in arbitrary units.

### Protein extraction and Immunoblotting

Two hundred milligrams of leaf material were mixed with 350 µL of extraction buffer (100 mM Tris-HCl, pH 7.5, 150 mM NaCl, 0.1% Nonidet P-40 and 1x Protease inhibitor cocktail (Sigma, P599)) and centrifuged for 15 min at 14,000 *g*. 4X Laemmli sample was added to 100 µL supernatant and boiled for 5 min. Equal amounts of supernatant were loaded on 10% polyacrylamide gels and blotted on a polyvinylidene difluoride (PVDF) membrane. α-Myc and α-rabbit horseradish peroxidase conjugate (Amersham, GE Healthcare) were used at dilutions 1:1,000 and 1:5,000, respectively. The reaction was developed for 1 min using a Luminata Crescendo Millipore immunoblotting detection system (Millipore, WBLUR0500).

For detection of phosphorylated forms of SnRK1 proteins, 10-day-old seedlings were collected and ground in liquid nitrogen and the proteins were extracted using the following extraction buffer: 25 mM Tris-HCl pH 7.8, 75 mM NaCl, 15 mM EGTA, 10 mM MgCl_2_, 10 mM B-glycerophosphate, 15 mM 4-Nitrophenylphosphate bis, 1 mM DTT, 1 mM NaF, 0.5 mM Na_3_VO_4_, 0.5 mM PMSF, 1% Protease inhibitor cocktail (Sigma, P599), 0.1% Tween-20. The protein extracts were centrifuged at 13,000 rpm and 4°C for 10 min and supernatants transferred to a new tube. The protein concentration was measured using Bradford Dye Reagent (Bio-Rad); equal amounts (15 µg) of total protein for each sample were separated by SDS-PAGE (10% acrylamide gel) and transferred to a PVDF membrane (Bio-Rad). The membrane was blocked in TBST buffer containing 5% (w/v) BSA and incubated with primary antibody and secondary antibody. Antibodies used for immunoblotting were as follows: *α*–Phospho-AMPKα (Thr175) (*α*-pT175) (1:1,000, Cell Signaling Technology), *α*-Kin10 (1:1,000, Agrisera), *α*-Kin11 (1:1,000, Agrisera), and *α*-Actin (1:10,000, Agrisera).

### Co-immunoprecipitation (Co-Ip)

For Co-Ip assays, total proteins from 7-day-old seedlings were extracted with no-salt lysis buffer (50 mM Tris, pH 8.0, 0.1% Nonidet P-40, and 1% Protease inhibitor cocktail [Sigma]) at a fresh weight:buffer volume ratio of 1 g:2 mL. After centrifugation at 6,000 g and 4°C for 5 min, 20 µL of α-GFP microbeads (Miltenyi Biotec) were added to the resultant supernatant and incubated for 1 h at 4°C on a rotating wheel. Subsequent washing and elution steps were performed according to the manufacturer (µMACS GFP Isolation Kit; Miltenyi Biotec). Immunoblot analysis was done essentially as described above, and immunoprecipitates from transgenic lines expressing free GFP were used as controls. GFP-TSN-interacting proteins and native TSN were detected by mouse α -GFP (monoclonal antibody JL-8; Clontech) and rabbit α-TSN antibodies at final dilutions of 1:1,000 and 1:5,000, respectively.

### Image Analysis

The image analysis was done using ImageJ v1.41 (NIH) software (http://rsb.info.nih.gov/ij/index.html). Co-localization analyses were performed as described previously (French et al., 2008) using Pearson (rp) and Sparman (rs) statistics.

### Bioinformatics

Functional annotation of Gene Ontology was performed using Panther (Mi et al., 2017). Information about subcellular localization for all proteins was performed using SUBA4 (Tanz et al., 2013). Prion-like domains were identified using the web application PLAAC (Lancaster et al., 2014). The selected parameters were as follow: minimum length for prion domains was 60 (Lcore =60); organism background was *Arabidopsis*; and the α parameter was 1 (α =1). The RNA-binding proteins were predicted by the RNApred tool (Kumar et al., 2011). The selected parameters were as follows: prediction approach was amino acid composition and threshold for the support vector machine (SVM) was 0.5. To build protein-protein interaction networks we used STRING database (Jensen et al., 2009). We used Cytoscape to visualize the resulting PPI dataset (Gagneur et al., 2006). Per-residue disorder content was evaluated by PONDR predictors, including PONDR-FIT (Xue et al., 2010) and PONDR-VSL2 (Peng et al., 2005). The intrinsic disorder propensities of TSN were evaluated according to the method described by Uversky et al. (2017) (Santamaria et al., 2017; Uversky, 2017). Disorder evaluations together with disorder-related functional information were retrieved from the D_2_P_2_ database (http://d2p2.pro/) (Oates et al., 2013). LLPS predisposition was evaluated using the PSPredictor tool (Sun. et al., 2019).

## ACKNOWLEDGEMENTS

This work was supported by grants from European Commission (MSCA-IF-ReSGulating-702473) and Ministerio de Economía y Competitividad (Juan de la Cierva-Incorporacion grant, IJCI-2016-30763) to E.G.-B, and from Knut and Alice Wallenberg Foundation, the Swedish Research Council, the Swedish Foundation for Strategic Research, and Crops for the Future Research Programme to P.V.B.

## AUTHORŚCONTRIBUTIONS

Conceptualization, E.G.-B.; Methodology, E.G.-B., and P.V.B.; Investigation, E.G-B., P.H.E., V.U., and K.D.; Writing – Original Draft, E.G.-B.; Writing – Review & Editing, E.G.-B., P.N.M., K.D., J.L.C. and P.V.B.; Funding Acquisition, E.G.-B., and P.V.B.

---

**Figure S1.**
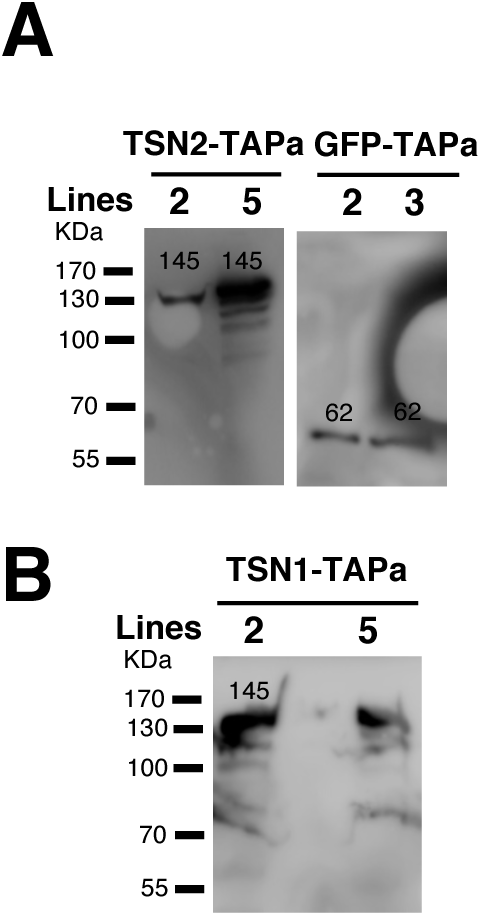
Transgenic *Arabidopsis* lines used in this study. *A*, Expression of TSN2-TAPa (167 KDa) and GFP-TAPa (83 KDa) and ***B***, TSN1-TAPa (170 KDa) in Col background confirmed by immunoblotting with α-Myc.

**Figure S2:**
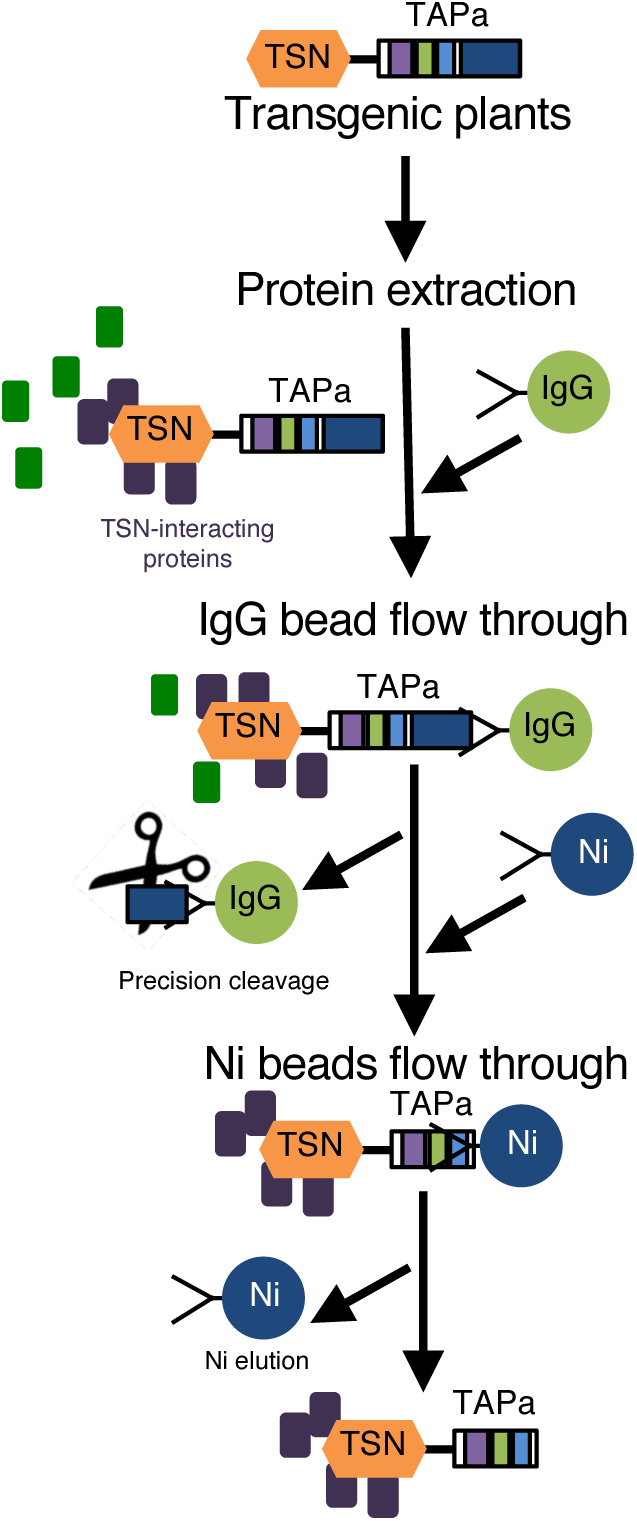
Schematic representation of the TAPa purification procedure. During the first affinity purification step, plant protein extracts are incubated with IgG beads followed by elution through the specific cleavage of TAPa tag with the low-temperature active rhinovirus 3C protease. At the second affinity purification step, IgG bead eluates are incubated with Ni beads followed by the elution of proteins from beads using imidazole-containing buffer.

**Figure S3.**
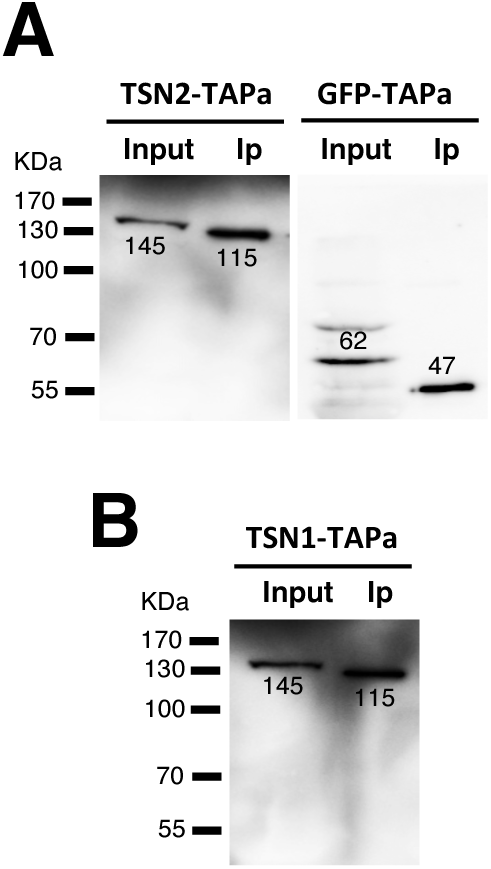
Immunoblotting of crude protein extracts (Input) and purified protein fractions (Ip) obtained during small-scale TAPa purification. TSN2-TAPa and GFP-TAPa (*A*) and TSN1-TAPa (*B*) fusion proteins were detected using α-Myc.

**Figure S4.**
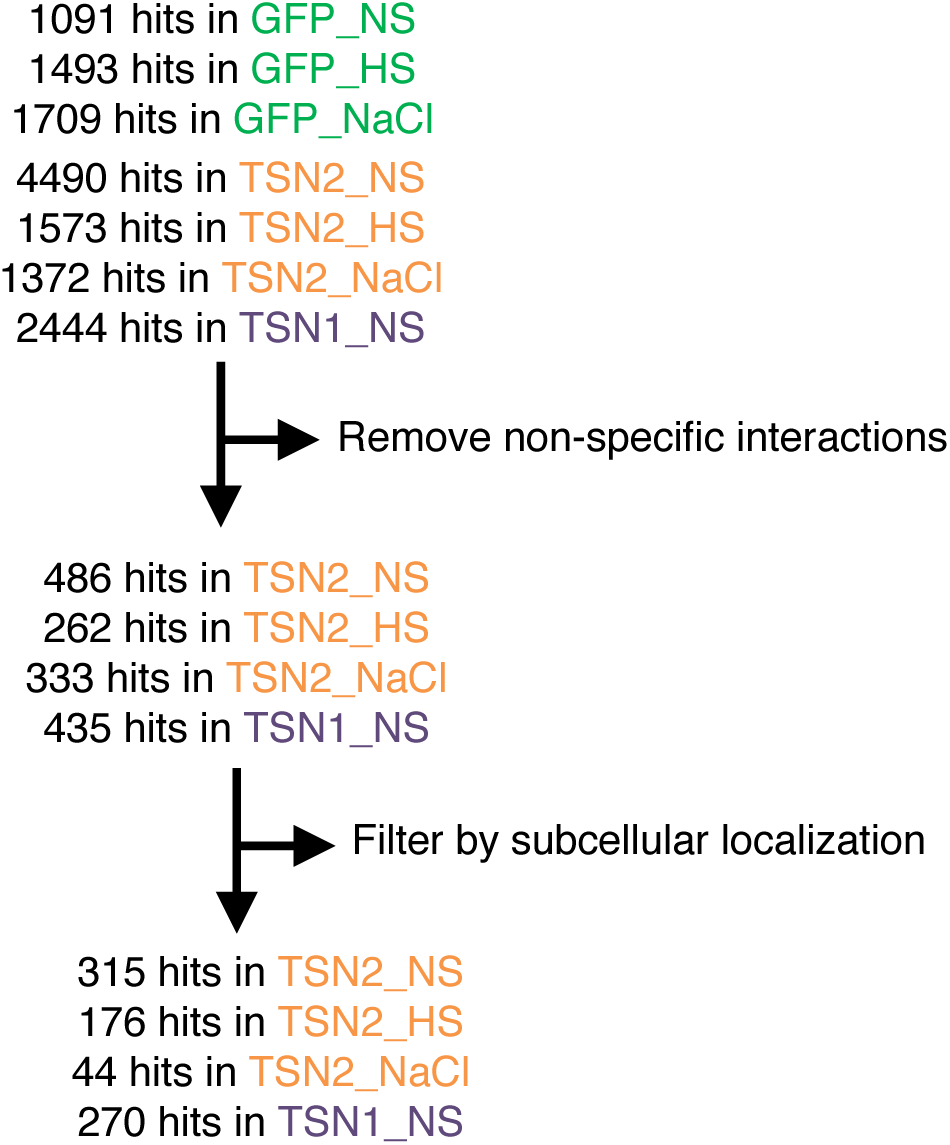
Flow chart of proteomics data generation.

**Figure S5.**
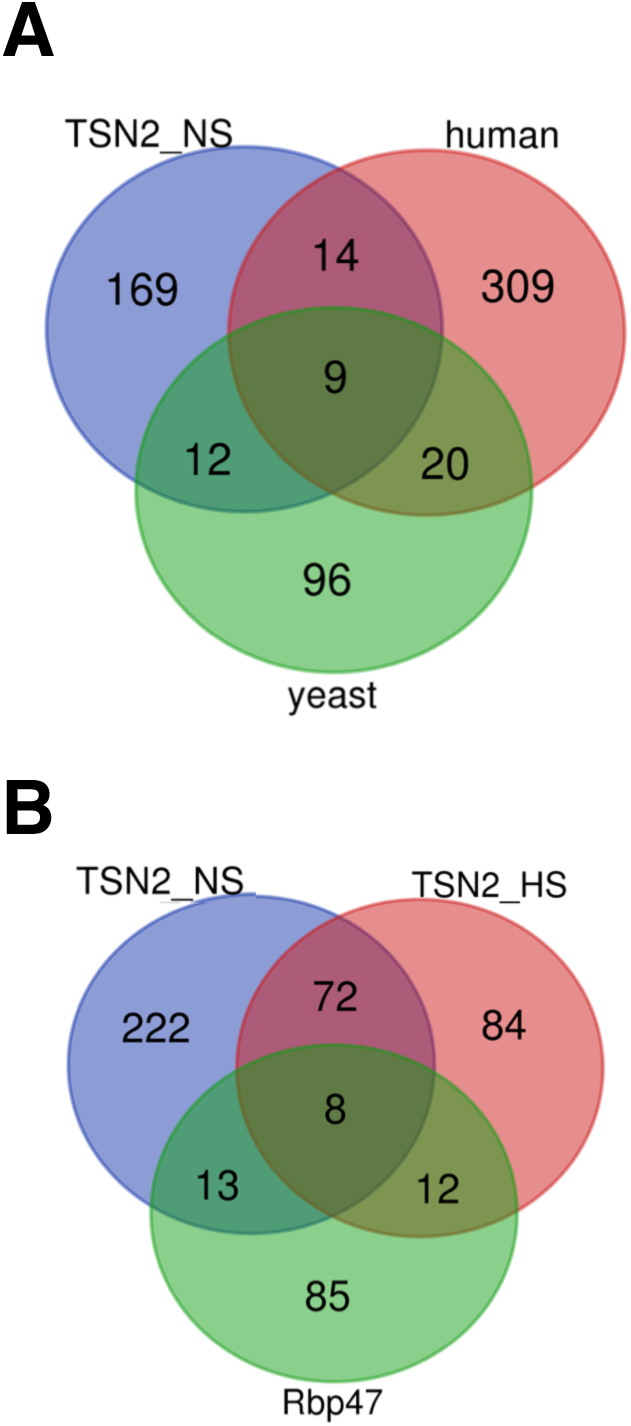
Venn diagrams. (***A*)** Comparison between mammalian and yeast SG proteomes with TSN2_NS interactome. (***B*)**, Comparison between TSN2_NS, TSN2_HS and Rbp47b proteomes.

**Figure S6.**
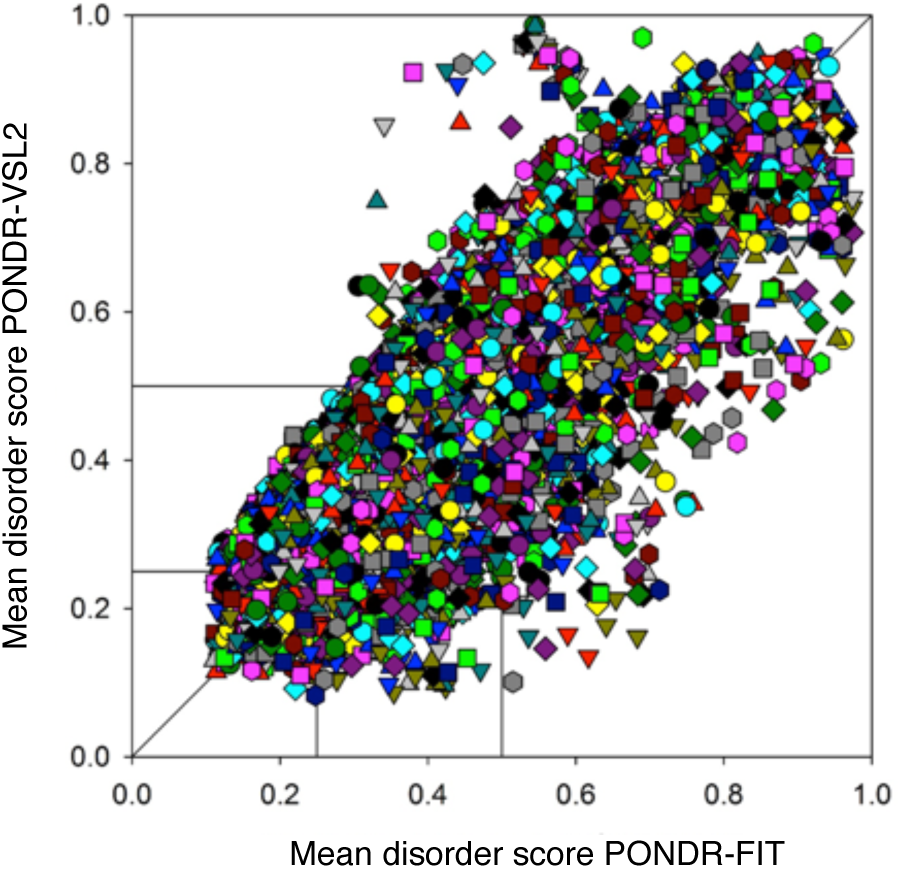
Analysis of the whole proteome of *Arabidopsis* for ID. Per-protein propensities for disorder (average of the corresponding per-residue propensities) evaluated by PONDR-FIT (*x*-axis) and by PONDR-VSL2 (y-axis).

**Figure S7.**
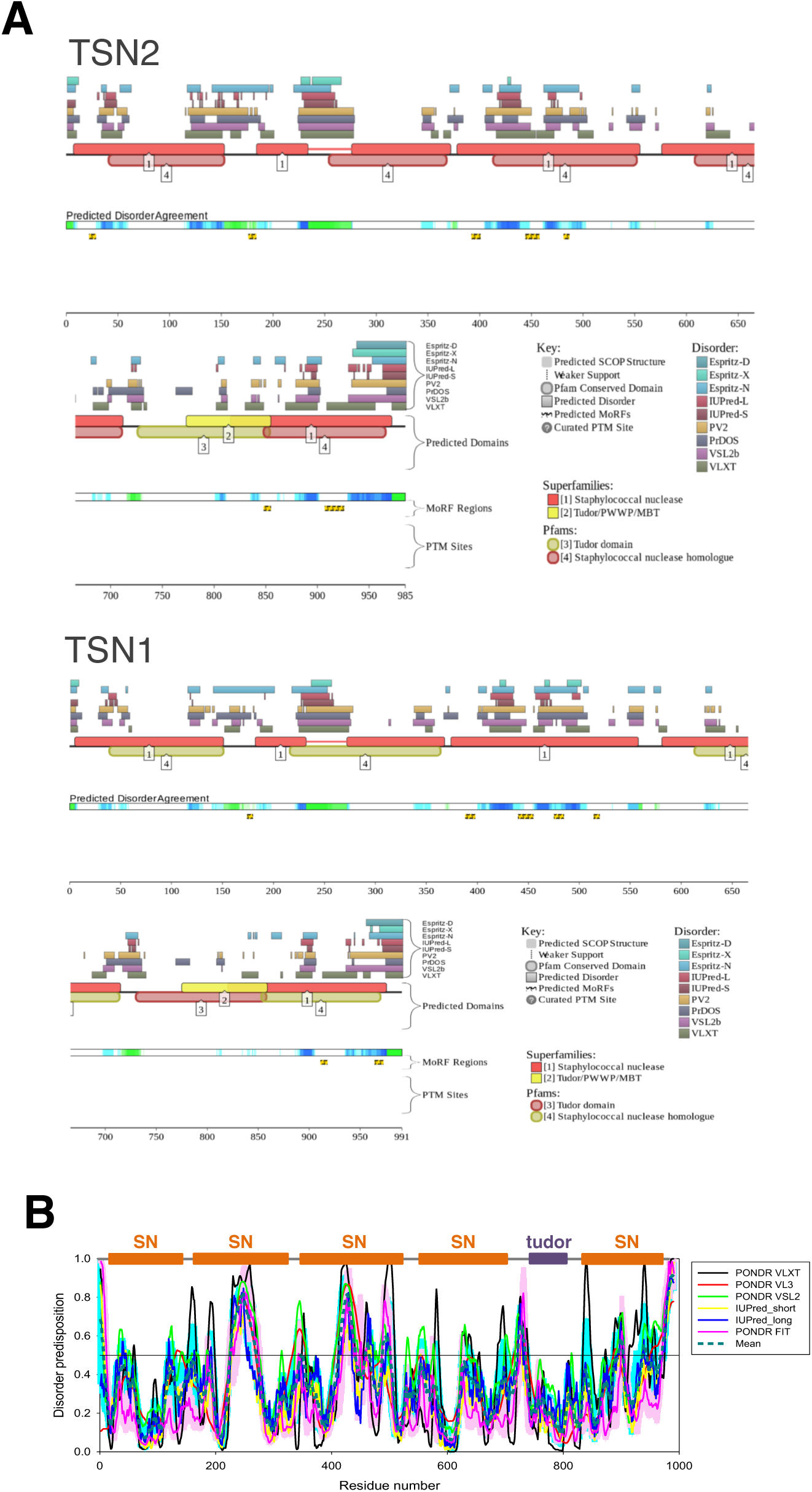
Prevalence and functionality of ID in the Atabidopsis TSN proteins. *A*, Evaluation of the functional ID propensity by the D2P2 database. In the corresponding plot, top nine colored bars represent the location of IDRs predicted by different disorder predictors (Espritz-D, Espritz-N, Espritz-X, IUPred-L, IUPred-S, PV2, PrDOS, PONDRs VSL2b, and PONDRs-VLXT, see keys for the corresponding color codes. Green/Blue-and-white bar in the middle of the plot shows the predicted disorder agreement between these nine predictors, with green/blue parts corresponding to IDRs by consensus. The yellow bar shows the location of the predicted disorder-based binding site (MoRF region). ***B***, Evaluation of the per-residue disorder propensity of TSN1 using six different disorder predictors, and a consensus disorder profile (based on mean values of six predictors). SN, staphylococcal nuclease domain.

**Figure S8.**
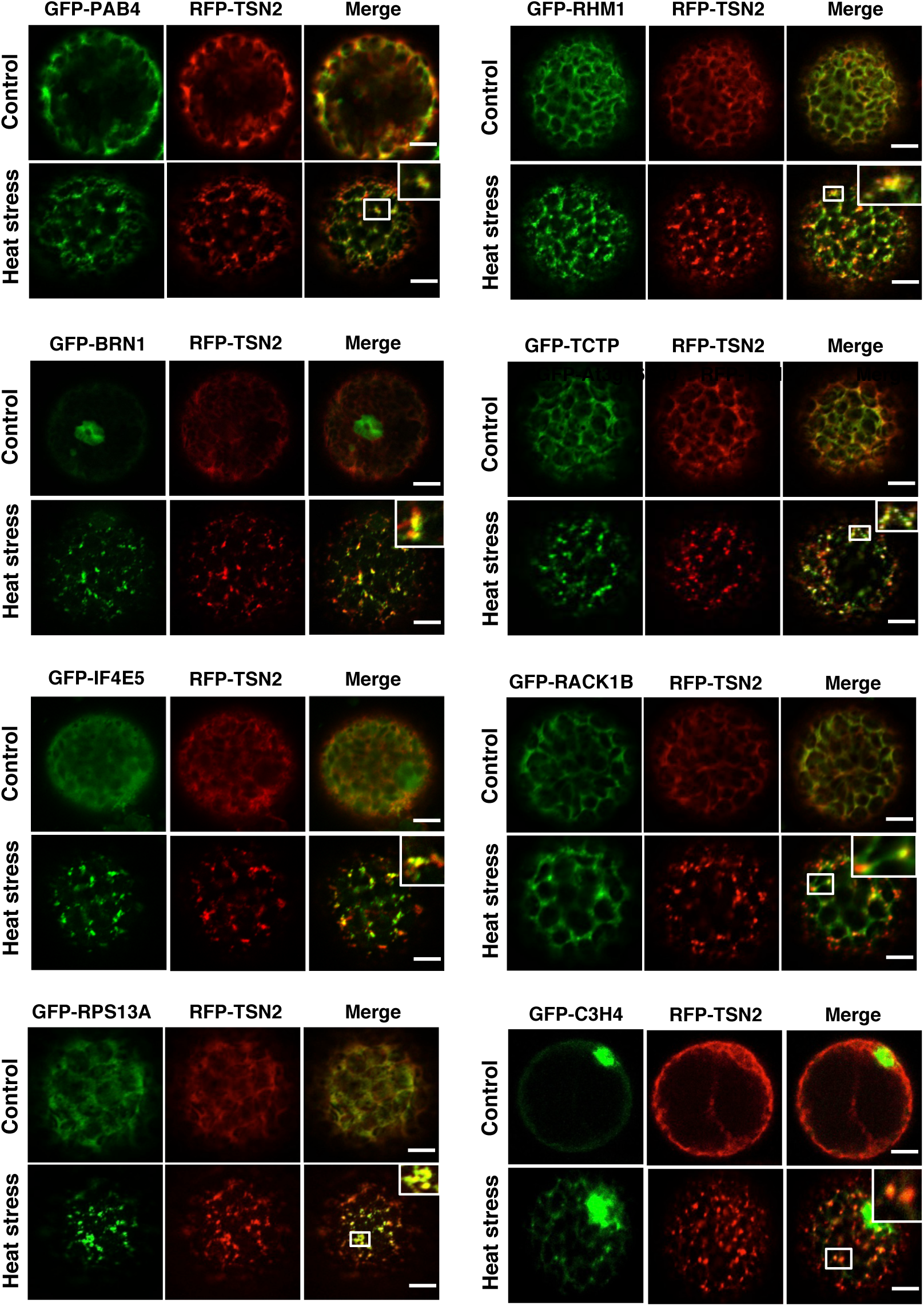
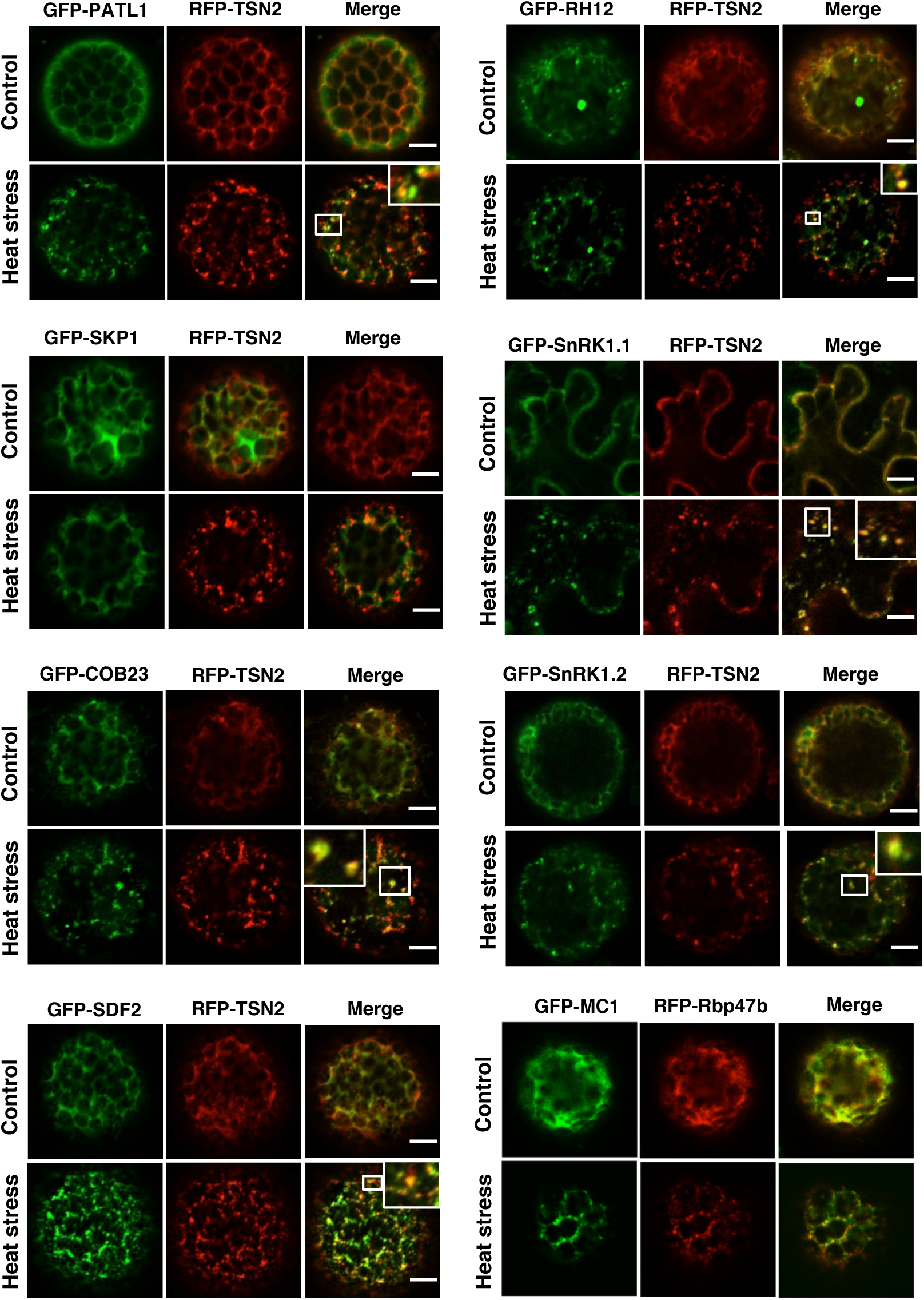
Co-localization of GFP-TSN2-interacting proteins (green) and RFP-TSN2 (red) quantified in Figure 4C. Insets show enlarged boxed areas. Scale bars = 5 µm.

**Figure S9.**
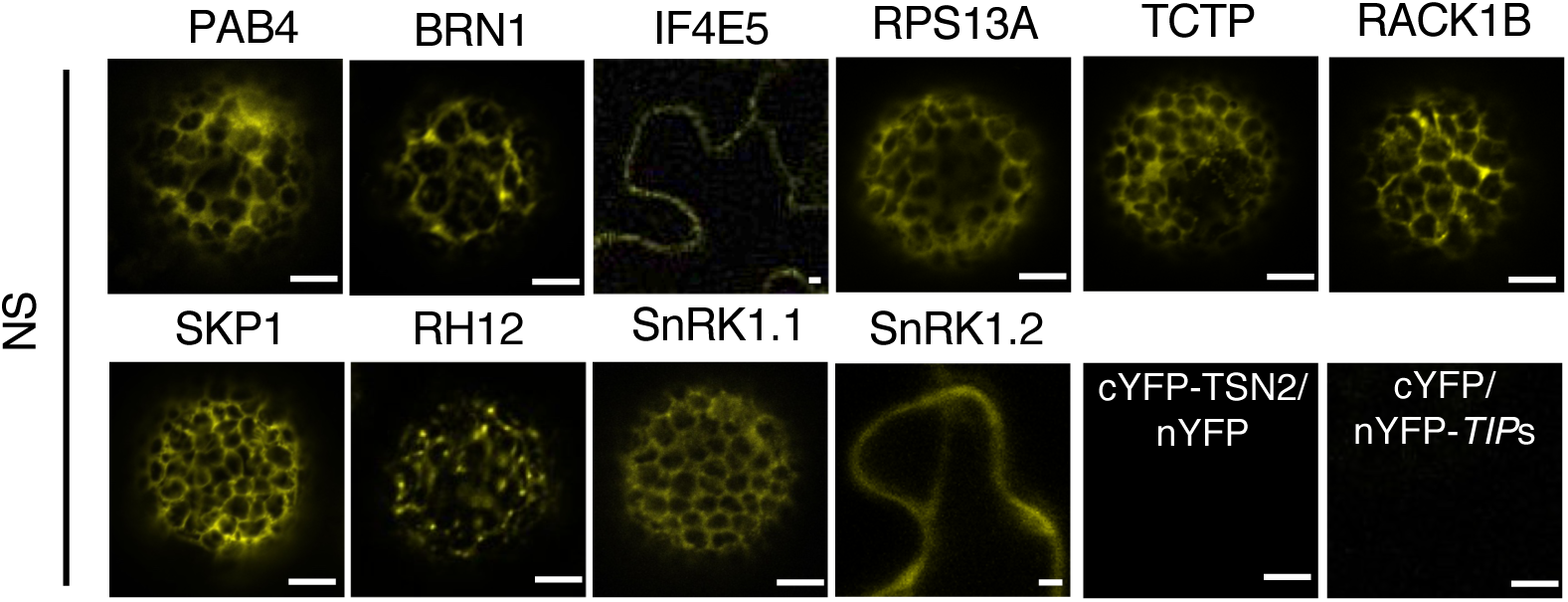
BiFC between TSN2 and TSN2-interacting proteins under no stress (NS) condition. BiFC between cYFP-TSN2 and nYFP-TSN-interacting proteins (TIPs) in *N. benthamiana* protoplasts incubated at 23°C. BiFC analysis of cYFP-TSN2 and nYFP-TIPs (only one representative example is shown) with empty vectors encoding nYFP and cYFP, respectively was used as a negative control. Scale bars = 5 µm.

**Figure S10.**
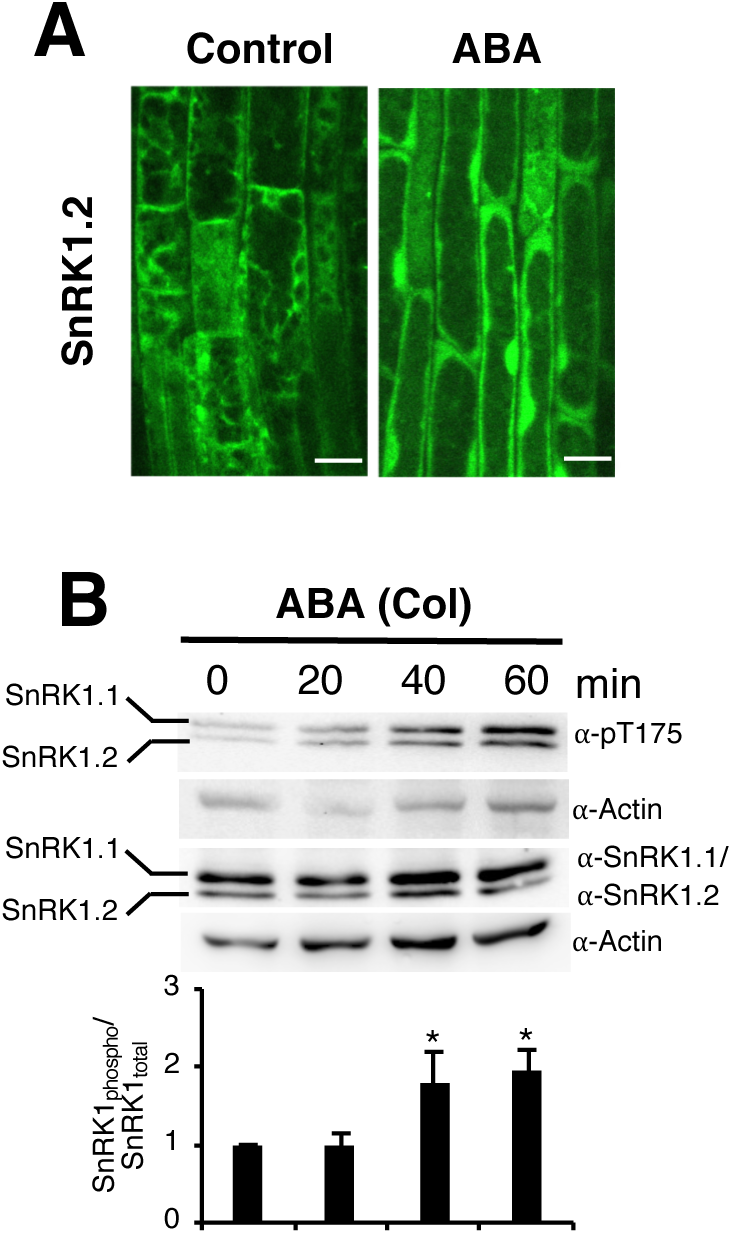
Activation of SnRK1 isoforms under ABA treatment. *A*, Localization of SnRK1.2 is shown in the root cells of 5-day-old Col seedlings expressing Pro35S:GFP-SnRK1.2. The seedlings were grown under control conditions or incubated for 40 min at 10 µM of ABA (ABA). Bars = 10 µm. ***B*,** Immunoblot analysis using α-SnRK1.1, α-SnRK1.2, α-p-T175 and α-Actin in ABA-stressed 10-day-old *Arabidopsis* Col seedlings for 0, 20, 40 and 60 min. ***C***, SnRK1 activity was determined as the ratio of phosphorylated to total SnRK1 protein. Three biological replicates were analysed for quantification. Asterisks denote significant difference; Student’s t-test, P < 0.05.

## REFERENCES

Abe S, Sakai M, Yagi K, Hagino T, Ochi K, Shibata K, Davies E (2003) A Tudor protein with multiple SNc domains from pea seedlings: cellular localization, partial characterization, sequence analysis, and phylogenetic relationships. J Exp Bot 54: 971–983

Arimoto K, Fukuda H, Imajoh-Ohmi S, Saito H, Takekawa M (2008) Formation of stress granules inhibits apoptosis by suppressing stress-responsive MAPK pathways. Nat Cell Biol 10: 1324–1332

Baena-Gonzalez E, Hanson J (2017) Shaping plant development through the SnRK1-TOR metabolic regulators. Curr Opin Plant Biol 35: 152–157

Baena-Gonzalez E, Rolland F, Thevelein JM, Sheen J (2007) A central integrator of transcription networks in plant stress and energy signalling. Nature 448: 938–942

Bogamuwa S, Jang JC (2013) The Arabidopsis tandem CCCH zinc finger proteins AtTZF4, 5 and 6 are involved in light-, abscisic acid- and gibberellic acid-mediated regulation of seed germination. Plant Cell Environ 36: 1507–1519

Buchan JR, Parker R (2009) Eukaryotic stress granules: the ins and outs of translation. Mol Cell 36: 932–941

Buchan JR, Yoon JH, Parker R (2011) Stress-specific composition, assembly and kinetics of stress granules in Saccharomyces cerevisiae. J Cell Sci 124: 228–239

Carroll B, Dunlop EA (2017) The lysosome: a crucial hub for AMPK and mTORC1 signalling. Biochem J 474: 1453–1466

Cary GA, Vinh DB, May P, Kuestner R, Dudley AM (2015) Proteomic Analysis of Dhh1 Complexes Reveals a Role for Hsp40 Chaperone Ydj1 in Yeast P-Body Assembly. G3 (Bethesda) 5: 2497–2511

Cazares-Apatiga J, Medina-Gomez C, Chavez-Munguia B, Calixto-Galvez M, Orozco E, Vazquez-Calzada C, Martinez-Higuera A, Rodriguez MA (2017) The Tudor Staphylococcal Nuclease Protein of Entamoeba histolytica Participates in Transcription Regulation and Stress Response. Front Cell Infect Microbiol 7: 52

Chantarachot. T, Sorenson. RS, Hummel. M, Ke. H, Kettenburg. AT, Chen. D, Aiyetiwa. K, Dehesh. K, Eulgem. T, Sieburth. LE, Bailey-Serres. J (2019) DHH1/DDX6-like RNA helicases maintain ephemeral half-lives of stress-response mRNAs associated with innate immunity and growth inhibitio. bioRxiv

Chittum HS, Lane WS, Carlson BA, Roller PP, Lung FD, Lee BJ, Hatfield DL (1998) Rabbit beta-globin is extended beyond its UGA stop codon by multiple suppressions and translational reading gaps. Biochemistry 37: 10866–10870

Clough SJ, Bent AF (1998) Floral dip: a simplified method for Agrobacterium-mediated transformation of Arabidopsis thaliana. Plant J 16: 735–743

Curtis MD, Grossniklaus U (2003) A gateway cloning vector set for high-throughput functional analysis of genes in planta. Plant Physiol 133: 462–469

dit Frey NF, Muller P, Jammes F, Kizis D, Leung J, Perrot-Rechenmann C, Bianchi MW (2010) The RNA binding protein Tudor-SN is essential for stress tolerance and stabilizes levels of stress-responsive mRNAs encoding secreted proteins in Arabidopsis. Plant Cell 22: 1575–1591

Franks TM, Lykke-Andersen J (2008) The control of mRNA decapping and P-body formation. Mol Cell 32: 605–615

French AP, Mills S, Swarup R, Bennett MJ, Pridmore TP (2008) Colocalization of fluorescent markers in confocal microscope images of plant cells. Nat Protoc 3: 619–628

Gagneur J, David L, Steinmetz LM (2006) Capturing cellular machines by systematic screens of protein complexes. Trends Microbiol 14: 336–339

Gao X, Fu X, Song J, Zhang Y, Cui X, Su C, Ge L, Shao J, Xin L, Saarikettu J, Mei M, Yang X, Wei M, Silvennoinen O, Yao Z, He J, Yang J (2015) Poly(A)(+) mRNA-binding protein Tudor-SN regulates stress granules aggregation dynamics. FEBS J 282: 874–890

Gao X, Shi X, Fu X, Ge L, Zhang Y, Su C, Yang X, Silvennoinen O, Yao Z, He J, Wei M, Yang J (2014) Human Tudor staphylococcal nuclease (Tudor-SN) protein modulates the kinetics of AGTR1-3’UTR granule formation. FEBS Lett 588: 2154–2161

Gilks N, Kedersha N, Ayodele M, Shen L, Stoecklin G, Dember LM, Anderson P (2004) Stress granule assembly is mediated by prion-like aggregation of TIA-1. Mol Biol Cell 15: 5383–5398

Gutierrez-Beltran E, Bozhkov PV, Moschou PN (2015) Tudor Staphylococcal Nuclease plays two antagonistic roles in RNA metabolism under stress. Plant Signal Behav 10: e1071005

Gutierrez-Beltran E, Denisenko TV, Zhivotovsky B, Bozhkov PV (2016) Tudor staphylococcal nuclease: biochemistry and functions. Cell Death Differ 23: 1739–1748

Gutierrez-Beltran E, Moschou PN, Smertenko AP, Bozhkov PV (2015) Tudor Staphylococcal Nuclease Links Formation of Stress Granules and Processing Bodies with mRNA Catabolism in Arabidopsis. Plant Cell

Gutierrez-Beltran E, Personat JM, de la Torre F, Del Pozo O (2017) A Universal Stress Protein Involved in Oxidative Stress Is a Phosphorylation Target for Protein Kinase CIPK6. Plant Physiol 173: 836–852

Heberle AM, Prentzell MT, van Eunen K, Bakker BM, Grellscheid SN, Thedieck K (2015) Molecular mechanisms of mTOR regulation by stress. Mol Cell Oncol 2: e970489

Hilliker A, Gao Z, Jankowsky E, Parker R (2011) The DEAD-box protein Ded1 modulates translation by the formation and resolution of an eIF4F-mRNA complex. Mol Cell 43: 962–972

Jain S, Wheeler JR, Walters RW, Agrawal A, Barsic A, Parker R (2016) ATPase-Modulated Stress Granules Contain a Diverse Proteome and Substructure. Cell 164: 487–498

Jensen LJ, Kuhn M, Stark M, Chaffron S, Creevey C, Muller J, Doerks T, Julien P, Roth A, Simonovic M, Bork P, von Mering C (2009) STRING 8--a global view on proteins and their functional interactions in 630 organisms. Nucleic Acids Res 37: D412–416

Jossier M, Bouly JP, Meimoun P, Arjmand A, Lessard P, Hawley S, Grahame Hardie D, Thomas M (2009) SnRK1 (SNF1-related kinase 1) has a central role in sugar and ABA signalling in Arabidopsis thaliana. Plant J 59: 316–328

Kedersha N, Ivanov P, Anderson P (2013) Stress granules and cell signaling: more than just a passing phase? Trends Biochem Sci 38: 494–506

Kedersha NL, Gupta M, Li W, Miller I, Anderson P (1999) RNA-binding proteins TIA-1 and TIAR link the phosphorylation of eIF-2 alpha to the assembly of mammalian stress granules. J Cell Biol 147: 1431–1442

Kosmacz M, Gorka M, Schmidt S, Luzarowski M, Moreno JC, Szlachetko J, Leniak E, Sokolowska EM, Sofroni K, Schnittger A, Skirycz A (2019) Protein and metabolite composition of Arabidopsis stress granules. New Phytol 222: 1420–1433

Kosmacz M, Luzarowski M, Kerber O, Leniak E, Gutierrez-Beltran E, Moreno JC, Gorka M, Szlachetko J, Veyel D, Graf A, Skirycz A (2018) Interaction of 2’,3’-cAMP with Rbp47b Plays a Role in Stress Granule Formation. Plant Physiol 177: 411–421

Krapp S, Greiner E, Amin B, Sonnewald U, Krenz B (2017) The stress granule component G3BP is a novel interaction partner for the nuclear shuttle proteins of the nanovirus pea necrotic yellow dwarf virus and geminivirus abutilon mosaic virus. Virus Res 227: 6–14

Kroschwald S, Maharana S, Mateju D, Malinovska L, Nuske E, Poser I, Richter D, Alberti S (2015) Promiscuous interactions and protein disaggregases determine the material state of stress-inducible RNP granules. Elife 4: e06807

Kumar M, Gromiha MM, Raghava GP (2011) SVM based prediction of RNA-binding proteins using binding residues and evolutionary information. J Mol Recognit 24: 303–313

Lancaster AK, Nutter-Upham A, Lindquist S, King OD (2014) PLAAC: a web and command-line application to identify proteins with prion-like amino acid composition. Bioinformatics 30: 2501–2502

Mahboubi H, Barise R, Stochaj U (2015) 5’-AMP-activated protein kinase alpha regulates stress granule biogenesis. Biochim Biophys Acta 1853: 1725–1737

Mahboubi H, Stochaj U (2017) Cytoplasmic stress granules: Dynamic modulators of cell signaling and disease. Biochim Biophys Acta 1863: 884–895

Maldonado-Bonilla LD (2014) Composition and function of P bodies in Arabidopsis thaliana. Front Plant Sci 5: 201

Markmiller S, Soltanieh S, Server KL, Mak R, Jin W, Fang MY, Luo EC, Krach F, Yang D, Sen A, Fulzele A, Wozniak JM, Gonzalez DJ, Kankel MW, Gao FB, Bennett EJ, Lecuyer E, Yeo GW (2018) Context-Dependent and Disease-Specific Diversity in Protein Interactions within Stress Granules. Cell 172: 590–604 e513

Martin K, Kopperud K, Chakrabarty R, Banerjee R, Brooks R, Goodin MM (2009) Transient expression in Nicotiana benthamiana fluorescent marker lines provides enhanced definition of protein localization, movement and interactions in planta. Plant J 59: 150–162

Martinez JP, Perez-Vilaro G, Muthukumar Y, Scheller N, Hirsch T, Diestel R, Steinmetz H, Jansen R, Frank R, Sasse F, Meyerhans A, Diez J (2013) Screening of small molecules affecting mammalian P-body assembly uncovers links with diverse intracellular processes and organelle physiology. RNA Biol 10: 1661–1669

Meng F, Na I, Kurgan L, Uversky VN (2015) Compartmentalization and Functionality of Nuclear Disorder: Intrinsic Disorder and Protein-Protein Interactions in Intra-Nuclear Compartments. Int J Mol Sci 17

Mi H, Huang X, Muruganujan A, Tang H, Mills C, Kang D, Thomas PD (2017) PANTHER version 11: expanded annotation data from Gene Ontology and Reactome pathways, and data analysis tool enhancements. Nucleic Acids Res 45: D183–D189

Molliex A, Temirov J, Lee J, Coughlin M, Kanagaraj AP, Kim HJ, Mittag T, Taylor JP (2015) Phase separation by low complexity domains promotes stress granule assembly and drives pathological fibrillization. Cell 163: 123–133

Nakamura S, Mano S, Tanaka Y, Ohnishi M, Nakamori C, Araki M, Niwa T, Nishimura M, Kaminaka H, Nakagawa T, Sato Y, Ishiguro S (2010) Gateway binary vectors with the bialaphos resistance gene, bar, as a selection marker for plant transformation. Biosci Biotechnol Biochem 74: 1315–1319

Nakaya A, Katayama T, Itoh M, Hiranuka K, Kawashima S, Moriya Y, Okuda S, Tanaka M, Tokimatsu T, Yamanishi Y, Yoshizawa AC, Kanehisa M, Goto S (2013) KEGG OC: a large-scale automatic construction of taxonomy-based ortholog clusters. Nucleic Acids Res 41: D353–357

Nukarinen E, Nagele T, Pedrotti L, Wurzinger B, Mair A, Landgraf R, Bornke F, Hanson J, Teige M, Baena-Gonzalez E, Droge-Laser W, Weckwerth W (2016) Quantitative phosphoproteomics reveals the role of the AMPK plant ortholog SnRK1 as a metabolic master regulator under energy deprivation. Sci Rep 6: 31697

Oates ME, Romero P, Ishida T, Ghalwash M, Mizianty MJ, Xue B, Dosztanyi Z, Uversky VN, Obradovic Z, Kurgan L, Dunker AK, Gough J (2013) D(2)P(2): database of disordered protein predictions. Nucleic Acids Res 41: D508–516

Ohn T, Kedersha N, Hickman T, Tisdale S, Anderson P (2008) A functional RNAi screen links O-GlcNAc modification of ribosomal proteins to stress granule and processing body assembly. Nat Cell Biol 10: 1224–1231

Peng K, Vucetic S, Radivojac P, Brown CJ, Dunker AK, Obradovic Z (2005) Optimizing long intrinsic disorder predictors with protein evolutionary information. J Bioinform Comput Biol 3: 35–60

Protter DS, Parker R (2016) Principles and Properties of Stress Granules. Trends Cell Biol 26: 668–679

Rayman JB, Karl KA, Kandel ER (2018) TIA-1 Self-Multimerization, Phase Separation, and Recruitment into Stress Granules Are Dynamically Regulated by Zn(2). Cell Rep 22: 59–71

Rodrigues A, Adamo M, Crozet P, Margalha L, Confraria A, Martinho C, Elias A, Rabissi A, Lumbreras V, Gonzalez-Guzman M, Antoni R, Rodriguez PL, Baena-Gonzalez E (2013) ABI1 and PP2CA phosphatases are negative regulators of Snf1-related protein kinase1 signaling in Arabidopsis. Plant Cell 25: 3871–3884

Rubio V, Shen Y, Saijo Y, Liu Y, Gusmaroli G, Dinesh-Kumar SP, Deng XW (2005) An alternative tandem affinity purification strategy applied to Arabidopsis protein complex isolation. Plant J 41: 767–778

Santamaria N, Alhothali M, Alfonso MH, Breydo L, Uversky VN (2017) Intrinsic disorder in proteins involved in amyotrophic lateral sclerosis. Cell Mol Life Sci 74: 1297–1318

Shaw RJ (2009) LKB1 and AMP-activated protein kinase control of mTOR signalling and growth. Acta Physiol (Oxf) 196: 65–80

Sorenson R, Bailey-Serres J (2014) Selective mRNA sequestration by OLIGOURIDYLATE-BINDING PROTEIN 1 contributes to translational control during hypoxia in Arabidopsis. Proc Natl Acad Sci U S A 111: 2373–2378

Sun. T, Li. Q, Xu. Y, Zhang. Z, Lai. L, Pei. J (2019) Prediction of liquid-liquid phase separation proteins using machine learning. bioRxiv

Takahara T, Maeda T (2012) Transient sequestration of TORC1 into stress granules during heat stress. Mol Cell 47: 242–252

Tanz SK, Castleden I, Hooper CM, Vacher M, Small I, Millar HA (2013) SUBA3: a database for integrating experimentation and prediction to define the SUBcellular location of proteins in Arabidopsis. Nucleic Acids Res 41: D1185–1191

Thomas MG, Loschi M, Desbats MA, Boccaccio GL (2011) RNA granules: the good, the bad and the ugly. Cell Signal 23: 324–334

Tsai NP, Wei LN (2010) RhoA/ROCK1 signaling regulates stress granule formation and apoptosis. Cell Signal 22: 668–675

Uversky VN (2017) How to Predict Disorder in a Protein of Interest. Methods Mol Biol 1484: 137–158

Weber C, Nover L, Fauth M (2008) Plant stress granules and mRNA processing bodies are distinct from heat stress granules. Plant J 56: 517–530

Weissbach R, Scadden AD (2012) Tudor-SN and ADAR1 are components of cytoplasmic stress granules. RNA 18: 462–471

Wheeler JR, Matheny T, Jain S, Abrisch R, Parker R (2016) Distinct stages in stress granule assembly and disassembly. Elife 5

Wippich F, Bodenmiller B, Trajkovska MG, Wanka S, Aebersold R, Pelkmans L (2013) Dual specificity kinase DYRK3 couples stress granule condensation/dissolution to mTORC1 signaling. Cell 152: 791–805

Wisniewski JR, Zougman A, Nagaraj N, Mann M (2009) Universal sample preparation method for proteome analysis. Nat Methods 6: 359–362

Wolozin B, Apicco D (2015) RNA binding proteins and the genesis of neurodegenerative diseases. Adv Exp Med Biol 822: 11–15

Wu FH, Shen SC, Lee LY, Lee SH, Chan MT, Lin CS (2009) Tape-Arabidopsis Sandwich - a simpler Arabidopsis protoplast isolation method. Plant Methods 5: 16

Xue B, Dunbrack RL, Williams RW, Dunker AK, Uversky VN (2010) PONDR-FIT: a meta-predictor of intrinsically disordered amino acids. Biochim Biophys Acta 1804: 996–1010

Yamada K, Lim J, Dale JM, Chen H, Shinn P, Palm CJ, Southwick AM, Wu HC, Kim C, Nguyen M, Pham P, Cheuk R, Karlin-Newmann G, Liu SX, Lam B, Sakano H, Wu T, Yu G, Miranda M, Quach HL, Tripp M, Chang CH, Lee JM, Toriumi M, Chan MM, Tang CC, Onodera CS, Deng JM, Akiyama K, Ansari Y, Arakawa T, Banh J, Banno F, Bowser L, Brooks S, Carninci P, Chao Q, Choy N, Enju A, Goldsmith AD, Gurjal M, Hansen NF, Hayashizaki Y, Johnson-Hopson C, Hsuan VW, Iida K, Karnes M, Khan S, Koesema E, Ishida J, Jiang PX, Jones T, Kawai J, Kamiya A, Meyers C, Nakajima M, Narusaka M, Seki M, Sakurai T, Satou M, Tamse R, Vaysberg M, Wallender EK, Wong C, Yamamura Y, Yuan S, Shinozaki K, Davis RW, Theologis A, Ecker JR (2003) Empirical analysis of transcriptional activity in the Arabidopsis genome. Science 302: 842–846

Yan C, Yan Z, Wang Y, Yan X, Han Y (2014) Tudor-SN, a component of stress granules, regulates growth under salt stress by modulating GA20ox3 mRNA levels in Arabidopsis. J Exp Bot

Yang X, Shen Y, Garre E, Hao X, Krumlinde D, Cvijovic M, Arens C, Nystrom T, Liu B, Sunnerhagen P (2014) Stress granule-defective mutants deregulate stress responsive transcripts. PLoS Genet 10: e1004763

Youn JY, Dunham WH, Hong SJ, Knight JDR, Bashkurov M, Chen GI, Bagci H, Rathod B, MacLeod G, Eng SWM, Angers S, Morris Q, Fabian M, Cote JF, Gingras AC (2018) High-Density Proximity Mapping Reveals the Subcellular Organization of mRNA-Associated Granules and Bodies. Mol Cell 69: 517–532 e511

Zhu L, Tatsuke T, Mon H, Li Z, Xu J, Lee JM, Kusakabe T (2013) Characterization of Tudor-sn-containing granules in the silkworm, Bombyx mori. Insect Biochem Mol Biol 43: 664–674

